# Phostensin Enables Lymphocyte Integrin Activation and Population of Peripheral Lymphoid Organs

**DOI:** 10.1101/2021.09.24.461584

**Authors:** Ho-Sup Lee, Hao Sun, Frédéric Lagarrigue, Jay W. Fox, Nicholas E. Sherman, Alexandre R. Gingras, Mark H. Ginsberg

## Abstract

Rap1 GTPase drives assembly of the Mig-10/RIAM/lamellipodin–Integrin–Talin (MIT) complex that enables integrin-dependent lymphocyte functions. Here we used tandem affinity tag-based proteomics to isolate and analyze the MIT complex and reveal that Phostensin (PTSN), a regulatory subunit of protein phosphatase 1, is a component of the complex. PTSN mediates de-phosphorylation of Rap1 thereby preserving the activity and membrane localization of Rap1 to stabilize the MIT complex. CRISPR/Cas9-induced deletion of *PPP1R18,* which encodes PTSN, markedly suppresses integrin activation in Jurkat human T cells. We generated apparently healthy *Ppp1r18^-/-^* mice that manifest lymphocytosis and reduced population of peripheral lymphoid tissues ascribable to defective activation of integrins α_L_β_2_ and α_4_β_7_. *Ppp1r18^-/-^* T cells exhibit reduced capacity to induce colitis in a murine adoptive transfer model. Thus, PTSN enables lymphocyte integrin-mediated functions by dephosphorylating Rap1 to stabilize the MIT complex. As a consequence, loss of PTSN ameliorates T cell-mediated colitis.

**SUMMARY:** Phostensin, a protein phosphatase 1 regulatory subunit, supports lymphocyte integrin-dependent functions by mediating dephosphorylation of Rap1 to stabilize the MIT complex thereby enabling the population of peripheral lymphoid organs and T cell-mediated colitis.

## INTRODUCTION

Integrin-mediated lymphocyte adhesion plays an essential role in lymphocyte development and their capacity to populate lymphoid organs and sites of inflammation. Furthermore, by playing a critical accessory role in the formation of immunological synapses, integrins facilitate processes such as cytotoxic killing and antigen presentation. Integrins α_L_β_2_ (LFA-1), α_4_β_1_ (VLA-4) and α_4_β_7_ (LPAM) are the lymphocyte integrins most commonly involved in these functions and each has served as a therapeutic target in autoimmune and inflammatory diseases(Hogg et al., 2011; Ley et al., 2016). Lymphocyte integrins are constitutively in a low-affinity state until agonist stimulation induces high-affinity, a process operationally defined as “integrin activation”(Luo et al., 2007; Takagi et al., 2002; Xiong et al., 2001). Integrin activation is essential to their capacity to mediate the lymphocyte functions enumerated above(Hynes, 2002; Ley et al., 2007).

Stimulation via agonists such as chemokines or cytokines initiates a spectrum of intracellular signaling pathways to trigger integrin activation. These pathways ultimately converge upon the binding of talin1 to the integrin β cytoplasmic tail(Kim et al., 2011; Lefort et al., 2012; Tadokoro et al., 2003; Ye et al., 2010), a final common step in integrin activation. The small GTPase Rap1 is a dominant hub in the signaling pathways that control the talin-integrin-interaction(Bos, 2005). In conventional T cells, Rap1 utilizes RIAM as an effector for integrin activation, whereas in CD4^+^Foxp3^+^ regulatory T cells, another MRL protein, Lamellipodin (LPD), makes a more important contribution(Sun et al., 2021). We previously showed that the MRL Protein-Integrin-Talin (MIT) complex forms the “sticky fingers” that drive cell protrusion and enable migratory pathfinding(Lagarrigue et al., 2015).

RIAM is abundant in hematopoietic cells. Unlike talin1 deletion, germline loss of RIAM in mice does not affect development, hemostasis, or platelet integrin function(Klapproth et al., 2015; Stritt et al., 2015; Su et al., 2015). However, RIAM plays an important role in the activation of β_2_ and β_7_ integrins in neutrophils, macrophages and T cells(Boussiotis et al., 1997; Klapproth et al., 2015; Medrano-Fernandez et al., 2013; Su et al., 2015; Sun et al., 2021). β_1_ and β_3_ integrin functions are less affected by the absence of RIAM in these leukocytes(Klapproth et al., 2015; Su et al., 2015). RIAM-deficient mice exhibit significant leukocytosis associated with leukocyte adhesion deficiency and impaired leukocyte extravasation. The apparently normal development and lack of bleeding in RIAM null mice, combined with their protection in models of autoimmune disease such as inflammatory bowel disease(Sun et al., 2021) or type I diabetes(Lagarrigue et al., 2017), suggests that targeting the regulation of the RIAM MIT complex could serve as a strategy to inhibit pathological inflammation or autoimmunity.

Here we analyzed the mechanisms that orchestrate the function of the MIT complex by using a tandem affinity purification tag to isolate the native complex and mass spectrometry to identify associated proteins. We focused our attention on Phostensin (PTSN), an actin-binding regulatory subunit of protein phosphatase 1 (PP1) encoded by the *PPP1R18* gene(Kao et al., 2007) because PTSN is mainly expressed in leukocytes(Lin et al., 2011) and shares similar tissue distribution with RIAM. Here, we report that PTSN is physically associated with MIT complex and that it stabilizes the complex by mediating the dephosphorylation of Rap1 thereby preserving Rap1 activity and enabling integrin activation. We generated viable and fertile *Ppp1r18^-/-^* mice that exhibited impaired population of peripheral lymphoid tissues and defective activation of T cell integrins α_L_β_2_ and α_4_β_7_. Because *Ppp1r18^-/-^* mice appear healthy yet have a defect in lymphocyte integrin function, we suggest that PTSN may be a therapeutic target in autoimmune and inflammatory diseases. In support of this idea, *Ppp1r18^-/-^* T cells manifested reduced capacity to induce colitis in an adoptive transfer model. In sum, we identify PTSN as a new regulator of lymphocyte integrin-mediated functions, show that it acts via stabilization of the RIAM MIT complex by dephosphorylating Rap1, and identify it as a potential therapeutic target in T cell-mediated diseases.

## METHODS

### Cell lines and plasmids

U2-OS, HEK293 and 293A cells were grown in DMEM supplemented with 10%(v/v) FBS, non-essential amino acids, 1x Penicillin/Streptomycin, and 2mM L-glutamine at 37°C in a 5% CO_2_ incubator. Jurkat T cells were from ATCC and cultured in RPMI supplemented with 10%(v/v) FBS, non-essential amino acids, 1x Penicillin/Streptomycin, and 2 mM L-glutamine. U2-OS stable cell lines expressing α_ΙΙb_-streptavidin(SBP)-β_3_ or α_ΙΙb_-SBPβ_3_(D119A) were generated by lentiviral infection of α_IIb_-SBP with β_3_ or α_IIb_-SBP with β_3_(D119A), respectively. Monoclonal cell populations were selected by flow cytometry.

Mammalian expression plasmids encoding full-length integrin α_IIb_-SBP and β_3_ (WT, wild-type) or (D119A) were cloned into pcDNA3.1(-). Myc-, Myc-mCherry- or DsRed-tagged human β- (aa 1-613) and α-PTSN (aa 249-613) were cloned into pcDNA3.1(-). Full-length RIAM cDNA was cloned into p3xFlag-CMV-7.1 (Sigma-Aldrich, St. Louis, MO). Constructs encoding EGFP-tagged full-length talin1 WT were previously described(Wegener et al., 2007). Coding sequences of β- and α- PTSN were cloned into pEGFP-C1 and p3xFlag-CMV-7.1 (Sigma-Aldrich). β- PTSN 4A mutation (residues K539, I540, S541 and F542 into four Ala) predicted to block binding to protein phosphatase 1 (PP1)(Kao et al., 2007) was produced by site-directed mutagenesis. Bicistronic constructs expressing Flag-RIAM together with either Myc-His-Rap1a WT or S180A mutant were generated into pcDNA3.1(-). Plasmid encoding EGFP-tagged PP1 catalytic subunit γ was obtained from Addgene (Addgene plasmid # 44225). Expression of PTSN was silenced in U2-OS and 293A cells by lentiviral transduction of a pLKO1 shRNA against both β- and α-PTSN (TRCN0000282572, Sigma-Aldrich). pLKO1 shRNA (SHC016V, Sigma-Aldrich) was used as a control.

### Antibodies and reagents

Antibodies against talin (8d4, Sigma-Aldrich) and Flag (M2) were from Sigma-Aldrich. Antibody against DsRed (sc-33354) was from Santa Cruz Biotechnology (Dallas, TX). Anti-integrin α_IIb_ (PMI-1), anti-β_3_ (Rb8257 and Rb8053)(Frelinger et al., 1990; Frelinger et al., 1988), activated anti-α_IIb_β_3_ (Pac1)(Shattil et al., 1985) were described previously. Polyclonal anti-EGFP antibody was produced in rabbits immunized with recombinant EGFP. Anti-Flag M2 affinity gel was purchased from Sigma-Aldrich, and Pierce Streptavidin Plus UltraLink Resin and Pierce Monomeric Avidin Agarose were from Thermo Fisher Scientific (Rockford, IL). 3xFlag peptide (MDYKDHDGDYKDHDIDYKDDDDK) was synthesized by United BioSystems Inc (Hemdon, VA). Fluorophore-conjugated antibodies against CD3 (17A2, 2C11), B220 (RA3-6B2), CD49d (9F10), CD11a (M17/4), CD29 (HMβ1-1), CD18 (M18/2) and β_7_ (FIB 504) were purchased from BioLegend (San Diego, CA). Secondary AlexaFluor-labelled antibodies were from Jackson ImmunoResearch (West Grove, PA). MojoSort mouse CD4 T cell isolation kit and SDF-1α were from BioLegend. Recombinant human VCAM-1-Fc, mouse ICAM-1-Fc and VCAM-1-Fc were from BioLegend. Recombinant mouse MAdCAM-1-Fc was from R&D Systems (Minneapolis, MN).

Antibody against β-PTSN was from Santa Cruz Biotechnology. To generate antibody against α-PTSN, mouse PTSN C-terminus (a.a. 429-594) was cloned into pETM-11 vector. His-PTSN fusion protein was solubilized from inclusion bodies with 6M urea and purified under denaturing conditions using His-Bind resin. The protein was dialyzed against PBS multiple times and used to raise polyclonal rabbit antisera (Abgent, San Diego, CA).

### Purification of MIT complex by split-tandem affinity purification

Sub-confluent U2-OS cells stably expressing integrin α_IIb_-SBP and β_3_(D119A) were transfected with plasmids encoding Flag-RIAM or encoding the indicated Flag-tagged bait for 24-48 hrs. After washing with PBS, cells were harvested and lysates were prepared in lysis buffer (50 mM Tris-HCl (pH7.4), 100 mM NaCl, 0.5% NP-40, 0.5 mM MgCl_2_, 0.5 mM CaCl_2_, 0.2 mM GMP-PNP, 10 mM N-Ethylmaleimide, 1 μM Calpeptin, PhosStop and protease inhibitor cocktails). Cell lysates were centrifuged at 22,000 *g* for 30 min. The supernatant was collected and precleared with anti-mouse IgG coupled to agarose beads (Sigma-Aldrich) for 2 hrs at 4°C. Cell lysates were then loaded into anti-Flag affinity column and incubated for 4 hrs at 4°C. Bound proteins were eluted with 200 μg/mL 3xFlag peptide after washing with lysis buffer. The eluate was subsequently loaded into monomeric avidin agarose or streptavidin plus ultralink column (Thermo Fisher Scientific) and incubated for 2 hrs at 4°C. The associated proteins were washed and eluted by 5 mM D-biotin or boiled in Laemmli buffer for SDS-polyacrylamide gel electrophoresis (SDS-PAGE) and immunoblotting.

### Identification of MIT-associated proteins

The gel piece was transferred to a siliconized tube and washed and unstained in 200 µL 50% methanol overnight. The gel pieces were dehydrated in acetonitrile, rehydrated in 30 µL of 10 mM dithiothreitol in 0.1 M ammonium bicarbonate and reduced at room temperature for 30 min. The DTT solution was removed and the sample alkylated in 30 µL of 50 mM iodoacetamide in 0.1 M ammonium bicarbonate at room temperature for 30 min. The reagent was removed, and the gel pieces dehydrated in 100 µL acetonitrile. The acetonitrile was removed, and the gel pieces rehydrated in 100 µL of 0.1 M ammonium bicarbonateand the pieces completely dried by vacuum centrifugation. The gel pieces were rehydrated in 20 ng/µL trypsin in 50mM ammonium bicarbonate and digested overnight at 37 °C and the peptides formed extracted from the polyacrylamide in two 30 µL aliquots of 50% acetonitrile/5% formic acid. These extracts were combined and evaporated to 15 µL for MS analysis performed as previously described(Goldfinger et al., 2007).

### Immunoprecipitation

Cells were harvested and lysed by NP-40 lysis buffer (50 mM Tris-HCl pH 7.4, 150 mM NaCl, 0.5% NP-40, 0.5 mM CaCl_2_, 0.5 mM MgCl_2_, 1 μM calpeptin, protease inhibitor cocktail) on ice for 15 min. The lysate was centrifuged at 14,000 rpm at 4°C for 15 min. Protein concentration of the lysate was determined by using a BCA assay (Thermo Fisher Scientific). The lysate was immunoprecipitated using designated primary antibodies with protein G resin (GenScript, Piscataway, NJ), or anti-Flag M2 affinity agarose gel at 4°C. Then, immune complexes were washed for extended times with the lysis buffer and separated by SDS-PAGE gels. Proteins on the gel were transferred to nitrocellulose membrane and probed with indicated primary and secondary antibodies.

### Flow cytometry

Cells isolated from mouse tissues were washed and resuspended in PBS containing 0.1% BSA and stained with conjugated antibody for 30 min at 4 °C. Then cells were washed twice before flow cytometry analysis using an Accuri C6 Plus (BD Biosciences, Franklin Lakes, NJ). Data were analysed using FlowJo software (BD Biosciences).

### Integrin activation assay

The activation state of α_IIb_(R995A)β_3_ was assessed by measuring the binding of the ligand mimetic anti-α_IIb_β_3_ monoclonal antibody PAC1 in two-color flow cytometric assays as described previously(O’Toole et al., 1994). PTSN expression in HEK293 stable cells expressing integrin α_IIb_(R995A)β_3_ was silenced by shRNA and the cells were transfected with shRNA-resistant EGFP-PTSN or EGFP vector as a transfection marker. After 24 hrs, cells were suspended and stained with PAC1. Then, cells were washed and analyzed by flow cytometry. Background binding was measured in presence of the α_IIb_β_3_ antagonist integrilin (10 μM) whereas maximum binding was quantified upon addition of the anti-LIBS6 monoclonal antibody(Frelinger et al., 1991) against α_IIb_β_3_ (2. μM). To measure the activation of α_4_β_1_(Rose et al., 2000), cells (5×10^5^) were resuspended in HBSS buffer (Mediatech, Manassas, VA) containing 5 μg/mL human VCAM-1-Fc for 30 min at 37°C. Afterward, cells were washed with 0.5 mL of HBSS buffer, then resuspended in the same buffer containing AlexaFluor647-conjugated anti-human antibody and incubated for 30 min on ice. Cells were washed twice prior to analysis by flow cytometry. Background binding was measured in presence of 5 mM EDTA whereas maximum binding was quantified upon addition of 1mM MnCl_2_.

### Flow chamber assay

A polystyrene Petri dish was coated with 10 μL of MAdCAM-1/Fc, VCAM-1/Fc or ICAM-1/Fc alone or with SDF-1α (2 μg/ml) in coating buffer (PBS, 10 mM NaHCO3, pH9.0) for 1 hr at 37°C followed by blocking with 2% BSA in coating buffer for 1 hr at 37°C. Cells were diluted 10^6^ cells/mL in HBSS (10 mM HEPES, 1 mM Ca^2+^/Mg^2+^) and immediately perfused through the flow chamber at a constant flow rate of 2 dyn/cm^2^. Adhesive interactions between the flowing cells and the coated substrates were assessed by manually tracking the motions of individual cells for 1 min as previously described(Sun et al., 2020; Sun et al., 2014). The motion of each adherent cell was monitored for 10 sec following the initial adhesion point, and two categories of cell adhesion were defined: rolling adhesion for cells with rolling motions for >10 sec with a velocity >1 μm/sec, whereas cells that remained adherent and stationary for >10 sec with a velocity <1 μm/sec were defined as arrested adherent cells. The total adherent cell number includes both rolling adherent and firm adherent cells.

### Mice

All animal experiments were approved by the Institutional Animal Care and Use Committee (IACUC) of the University of California, San Diego, and were conducted in accordance with federal regulations as well as institutional guidelines and regulations on animal studies. All mice were housed in specific pathogen-free conditions. C57BL/6J (CD45.1), C57BL/6J (CD45.2), and *Rag1^-/-^* mice were from The Jackson Laboratory (Bar Harbor, ME). For experiments, 8-12-week-old mice were used. All injections of cells were performed during the light cycle. All experiments were performed by comparing mutant mice with littermate controls. Mononuclear cells were isolated from mesenteric lymph node (MLN), Peyer’s patch (PP), peripheral lymph node (PLN), spleen (SP) and colonic lamina propria as previously described. *Ppp1r18^-/-^* mice were generated using a CRISPR/Cas9 approach at the University of California Irvine Transgenic Mouse Facility. The complex Cas9 mRNA/sgRNA/tracRNA (3μM) was injected into the pronuclei of C57BL/6N embryos. Surviving embryos were implanted into ICR pseudo-pregnant females and pregnancies went to full term. Tissue biopsies for genomic DNA were taken from pups between 7-10 days. Mice were genotyped by PCR using forward primers 5’-ggacgacctgggacatagataca-3’ and reverse primers 5’- ttttcacacgccttcacaggta-3’ followed by Sanger sequencing using forward primers 5’- tccagacagcaggaagaggaag-3’ and reverse primers 5’- tctacccagtcaggcatggt-3’.

Founders were selected and backcrossed to the C57BL/6J strain (>5 times) to establish the *Ppp1r18^-/-^* strain. To generate the *Ppp1r18β^-/-^* mice, a similar CRISPR/Cas9 approach was used to inject a complexed Cas9 mRNA/sgRNA (3 μM) into the pronuclei of C57BL/6NHsd embryos at the University of California San Diego Transgenic Mouse Facility. One homozygous pup harbouring a 10 base pair deletion resulting in the appearance of a downstream premature stop codon was selected and backcrossed to the C57BL/6J strain (>5 times) to establish the *Ppp1r18β^-/-^* strain, which only lacks the β form of PTSN. Mice were genotyped by PCR using forward primers 5’-ctgagcagagacccactgaaag-3’ and reverse primers 5’- ggatctggcttctgagtttgtgta-3’ and amplicons were ran in 6% TBE gels (Life Technologies, Carlsbad, CA).

### Mouse colitis models

The adoptive T cell transfer model was described before(Sun et al., 2020). Briefly, 8-10-week-old mice were used. 5×10^5^ CD4^+^CD25^-^CD45RB^high^ conventional T cells from WT or *Ppp1r18^-/-^* mice were injected intraperitoneally into *Rag1^-/-^* mice(Mombaerts et al., 1992). All comparisons were made between littermates. Mouse body weight was measured every day and values are shown as a percentage of the original weight. During the duration of the experiment, we assessed the clinical progression of colitis by daily blinded scoring a disease activity index (DAI) by 2 independent investigators. The DAI is the combined score of body weight loss, stool consistency, and rectal bleeding and prolapse as follow: 1) Weight loss: 0 (no loss), 1 (1-5%), 2 (5-10%), 3 (10-20%), 4 (>20%); 2) Stool consistency: 0 (normal),1 (soft), 2 (very soft), 3 (diarrhea); 3) Rectal bleeding: 0 (none), 1 (red), 2 (dark red), 3 (gross bleeding); 4) Rectal prolapse: 0 (none), 1 (signs of prolapse), 2 (clear prolapse), 3 (extensive prolapse). Mice were sacrificed at week 15.

### Histology

Formalin-fixed, paraffin-embedded Swiss-rolled colon sections of 4-mm thickness were mounted on glass slides and followed by hematoxylin and eosin staining or periodic acid–schiff staining. Images were acquired with a Nanozoomer Slide Scanner (Hamamatsu, Hamamatsu City, Japan). Blinded histological scoring was performed by 2 investigators based on the method described previously(Erben et al., 2014) and total scoring range is 0-12.

### Blood counts

Peripheral blood was collected from the retro-orbital plexus and transferred to tubes containing K+.EDTA. Cell counts were performed using a Hemavet 950FS Hematology System programmed with mouse-specific settings (Drew Scientific. Miami Lakes, FL). All samples were tested in duplicate, and the mean for each animal was plotted.

### Real-time quantitative PCR analyses

Total RNA was isolated from colon using tissue homogenizer (JXFSTPRP-24, ThunderSci, Shanghai, China) and TRIzol reagent according to the manufacturer’s protocol (Thermo Fisher Scientific). For gene expression analysis, single-stranded cDNA was produced from 10 μg total RNA of colon using SuperScript III First-Strand synthesis and oligo-dT primers according to the manufacturer’s protocol (Thermo Fisher Scientific). Kappa SybrFast qPCR kit (Kapa Biosystems, Wilmington, MA) and thermal cycler (CFX96 Real-Time System; Bio-Rad, Hercules, CA) were used to determine the relative levels of the genes analysed (primer sequences are shown in Table 1) according to the manufacturer’s protocol. The 2^-ΔΔCT^ method was used for analysis, and data were normalized to GAPDH. Control values (*Rag1*^-/-^ mice injected with PBS) were set to 1 for comparisons.

### Statistics

Statistical significance was assayed by a two-tailed *t*-test for single comparisons. ANOVA with a Tukey post hoc test was used to assay statistical significance for multiple comparisons. A p value <0.05 was considered significant.

**Supplementary Material:** Supplementary Table 1 contains the proteomic analysis of the purified MIT complex, Supplementary Figures 1-3.

## RESULTS

### PTSN is a component of the MIT complex

To isolate an MIT complex, we devised a split tandem affinity purification method that isolates the native complex formed prior to ligand engagement (Fig. 1A). We transfected Flag-RIAM into U2-OS cells stably expressing integrin α_IIb_-streptavidin-binding peptide (SBP) and β_3_(D119A) to form ligand binding-defective α_IIb_(SBP)β_3_(D119A). RIAM and associated proteins were isolated by anti-Flag affinity chromatography of cell lysates and the native proteins were eluted with 200 μM Flag peptide. The eluate was then passed through a streptavidin column to bind the α_IIb_(SBP) and the native MIT complex was eluted with 5 mM biotin. In five experiments an average of 100 µg of complex was isolated from 2×10^7^ cells. The eluate of the streptavidin column was fractionated on SDS-PAGE and major protein components were visualized by immunoblotting with antibodies against talin, integrin, or RIAM (right, Fig. 1B). In a proteomic analysis (Supplementary Table 1), we noted that PTSN was a prominent component of the isolated complex. PTSN, encoded by *PPP1R18*, is an actin binding regulatory subunit of protein phosphatase 1 (PP1). PTSN shares these biochemical functions with Phactr4, which modulates integrin signaling and cofilin activity to coordinate enteric neural crest cell migration(Zhang et al., 2012). PTSN was also present in the LPD MIT complex (Fig. 1C). Importantly, PTSN was absent from the eluates of both Flag and streptavidin columns in the absence of Flag-RIAM (Fig. 1D).

**Figure 1.**
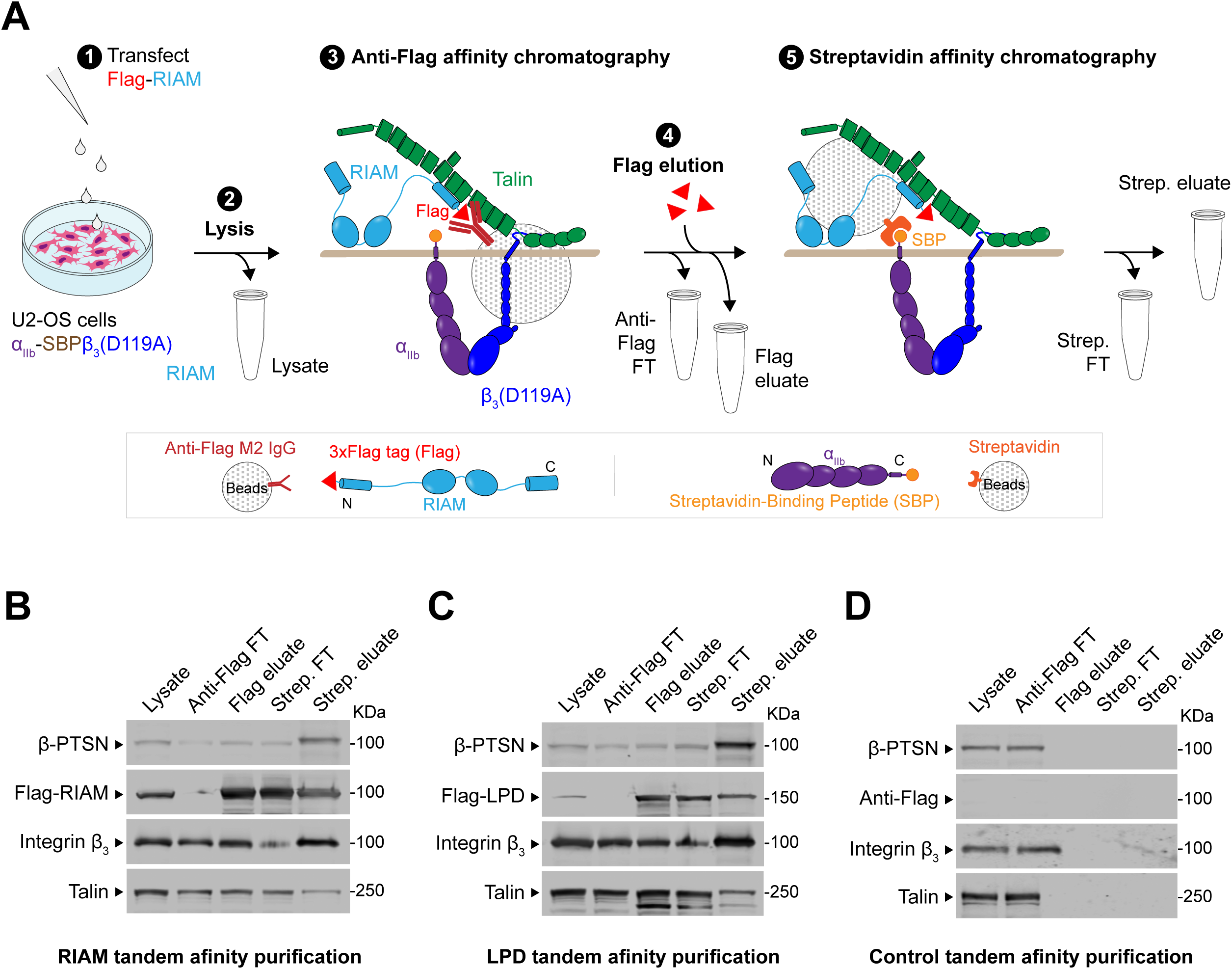
PTSN associates with the MIT complex. **(A)** Schematic of a split tandem affinity purification method to isolate the proteins associated with the MIT complex. **1:** U2-OS cells stably expressing integrin αIIb-streptavidin-binding peptide (SBP) and β_3_(D119A) to form ligand binding-defective α_IIb_(SBP)β_3_(D119A) were transfected with a plasmid encoding **(B)** Flag-tagged RIAM, **(C)** Flag-tagged LPD or **(D)** No insert. **2:** Cell lysis. **3:** Anti-Flag affinity chromatography of cell lysates. **4:** Native proteins were eluted with 200μM Flag peptide. **5:** The eluate was then passed through a streptavidin column. Bound MIT complexes were washed, eluted with 5 mM biotin and separated by SDS-PAGE. **(B-D)** Both RIAM- and LPD- MIT complexes contain PTSN. All fractions including whole cell lysate (0.5%), anti-Flag flow-through (FT, 0.5%), Flag eluate (10%), streptavidin flow-through (FT, 10%) and streptavidin eluate (50%) were analyzed by immunoblotting. Results are representative of 3 independent experiments.

### Both isoforms of PTSN regulate integrin activation

To examine the role of PTSN in integrin function, we used 293 cells constitutively expressing a recombinant talin-dependent(Tadokoro et al., 2003) activated integrin α_IIb_(R995A)β_3_. Silencing PTSN inhibited activation and co-expressing shRNA-resistant PTSN rescued activation of this integrin (Fig. 2A,B). PTSN has a short and long isoform, termed α and β respectively(Lin et al., 2014), and both contain actin and PP1 binding domains (Fig. 2C). Either isoform associated with RIAM (Fig. 2D) and could rescue integrin activation (Fig.2E).

**Figure 2.**
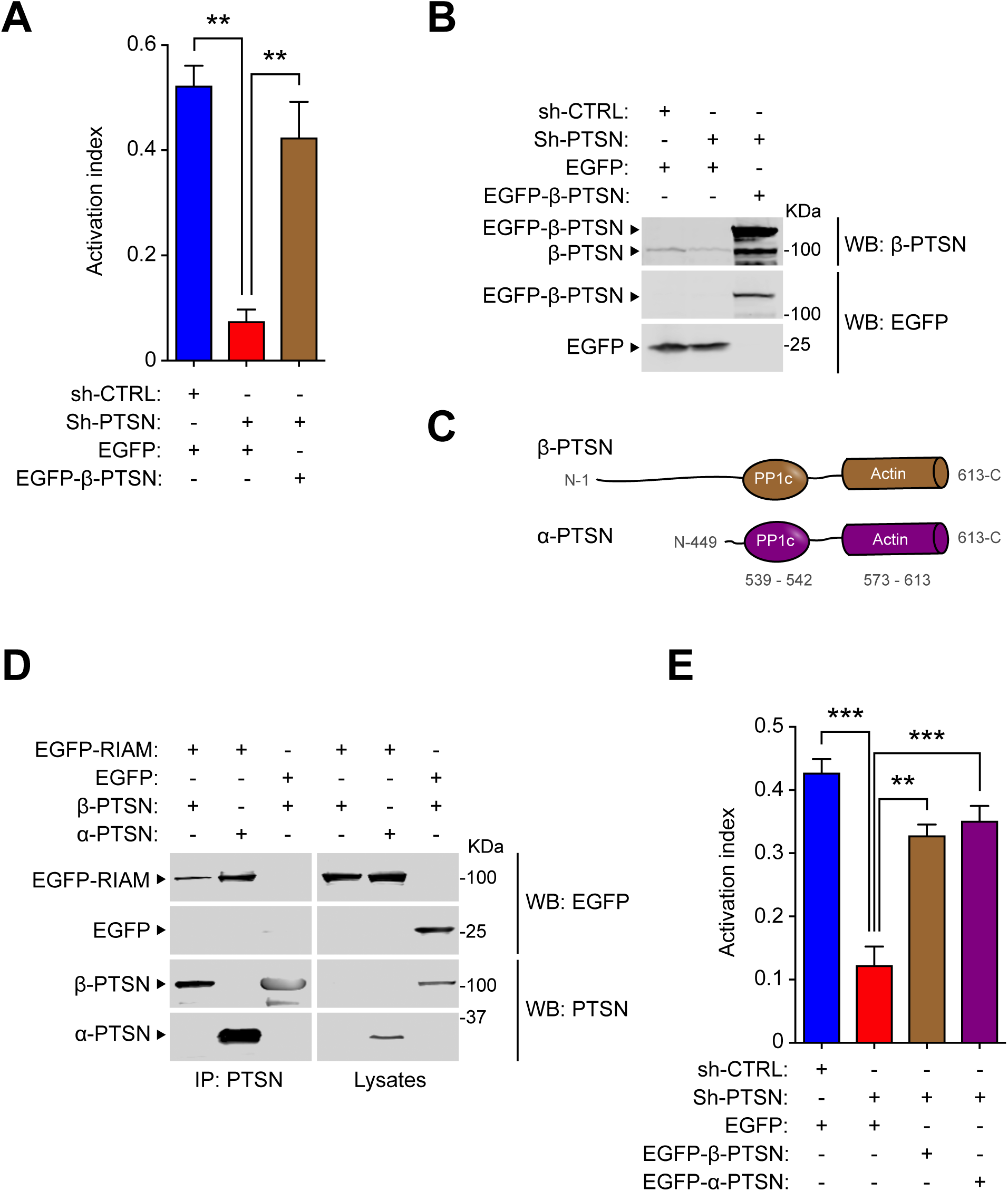
PTSN regulates integrin activation. **(A-B)** Silencing of PTSN reduces integrin activation. 293A cells that express constitutively active α_IIb_(R995A)β_3_ were transduced with a lentivirus encoding a shRNA against PTSN. A scrambled shRNA was used as a negative control. Cells were then transfected with a plasmid encoding either EGFP-tagged shRNA resistant PTSN or EGFP alone. **(A)** Integrin activation was measured by flow cytometry using the monoclonal antibody PAC1 that specifically recognizes the activated form of α_IIb_β_3_. The activation index was calculated as (*F* - *Fo*)/(*Fm* - *Fo*), in which *F* is the geometric mean fluorescence intensity (MFI) of PAC1 binding; *Fo* is the MFI in presence of the competitive inhibitor integrilin; and *Fm* is the MFI upon addition of the integrin-activating anti-LIBS6 antibody. Bar graphs represent mean ± SEM (n = 3 independent experiments). One-way ANOVA with Tukey post-test; ** p<0.01. **(B)** The expression of PTSN was confirmed by immunoblotting. **(C)** Domain organization of α- (short) and β-PTSN (long) isoforms. **(D)** PTSN α interacts with RIAM. HEK293 cells were transfected with plasmids encoding Flag-tagged α- or β-PTSN in combination with vectors encoding EGFP-RIAM or EGFP alone prior to anti-Flag affinity chromatography. **(E)** α-PTSN restores integrin activation in PTSN knockdown cells. The expression of PTSN was silenced in α_IIb_(R995A)β_3_ expressing 293A cells and the activation index was determined as in (A). Bar graphs represent mean ± SEM (n = 3 independent experiments). One-way ANOVA with Tukey post-test; ** p<0.01, *** p<0.001.

Analysis using the human proteome map (http://www.humanproteomemap.org/batch.php) confirmed that both RIAM and PTSN are particularly well expressed in lymphoid cells (Fig. 3A). We therefore used CRISPR/Cas9 mutagenesis to inactivate *PPP1R18* in the Jurkat T cell line and observed a marked reduction in basal and phorbol myristate acetate (PMA)- stimulated activation of an endogenous integrin (Fig. 3B,C). In both cases activation was rescued by re-expressing either α- or β-PTSN. Thus, both isoforms of PTSN regulate integrin activation in a model system and a human T cell line.

**Figure 3.**
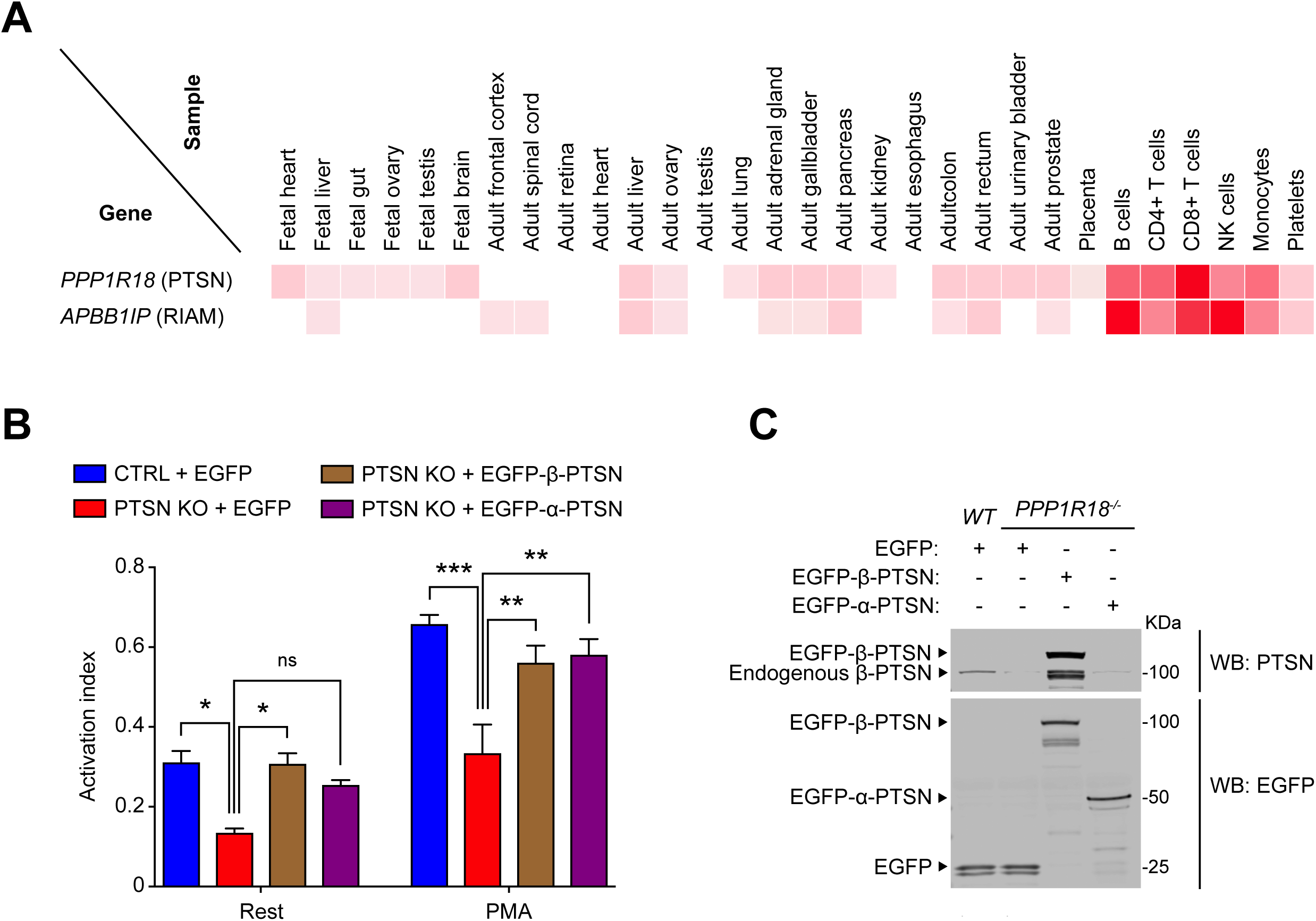
PTSN enables integrin activation in human T cell line. **(A)** Relative expression of PTSN and RIAM across various human tissues. Adapted from the Human Proteome Map portal (www.humanproteomemap.org). **(B)** Knockout of PTSN in Jurkat T cells reduces integrin activation. *PPP1R18* gene encoding PTSN was knocked-out in Jurkat T cells using CRISPR/Cas9 and soluble VCAM-1 binding was assessed by flow cytometry. Cellular stimulation was achieved using 25nM PMA. The activation index was calculated as (*F* - *Fo*)/(*Fm* - *Fo*), in which *F* is the mean fluorescence intensity (MFI) of VCAM-1 binding, *Fo* in presence of EDTA, and *Fm* upon addition of MnCl_2_. Bar graphs represent mean ± SEM (n = 3 independent experiments). Two-way ANOVA with Tukey post-test; ns not significant, * p<0.05, ** p<0.01, *** p<0.001. **(C)** The expression of PTSN was confirmed by immunoblotting. Results are representative of 3 independent experiments.

### PTSN regulates the assembly of the MIT complex

To investigate how PTSN regulates integrin activation, we tested the role of PTSN in the assembly of the RIAM MIT complex. We silenced PTSN expression in cells expressing recombinant Flag-RIAM and integrin α_IIb_β_3_ and purified Flag-RIAM by immuno-affinity chromatography (Fig. 4A). Silencing PTSN dramatically reduced the association of both integrin β_3_ and talin with RIAM, indicating disassembly of the MIT complex. The complex was assembled when shRNA-resistant β-PTSN was co-expressed with the PTSN shRNA (Fig. 4 B-E). Thus, PTSN regulates the assembly of the MIT complex.

**Figure 4.**
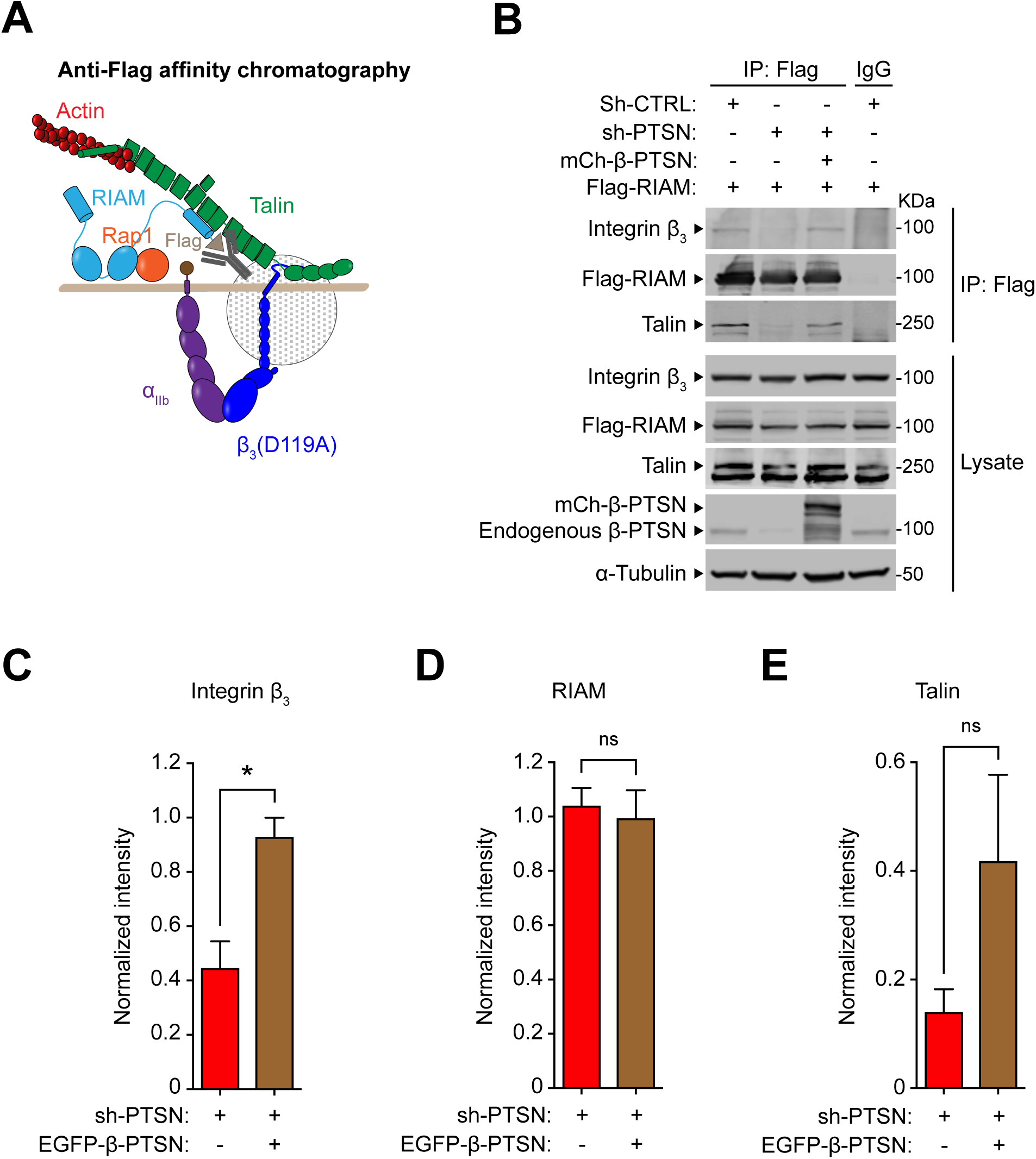
PTSN enables the assembly of the RIAM-MIT complex. **(A-B)** U2-OS cells that stably express integrin α_IIb_-SBP(streptavidin-binding peptide) and β_3_(D119A) to form ligand binding-defective α_IIb_(SBP)β_3_(D119A) were transduced with a lentivirus encoding a shRNA against PTSN prior to transfection with mCherry-tagged β-PTSN in combination with Flag-RIAM. Flag-tagged RIAM was captured by **(A)** anti-Flag immunoprecipitation and **(B)** associated proteins were analyzed by immunoblotting. **(C-E)** Integrated density values for immunoreactive bands corresponding to integrin β_3_ **(C)**, RIAM **(D)** or talin **(E)**. Bar graphs represent mean ± SEM (n = 3 independent experiments) normalized to control condition (sh-scrambled). Two-tailed *t*-test; ns not significant, * p<0.05.

Activated Rap1 drives the formation of the MIT complex and results integrin activation, leading us to examine the potential role of PTSN in regulating Rap1. Protein kinase A (PKA) phosphorylation of Rap1 inhibits Rap1 activity by decreasing GTP loading and disrupting membrane localization(Takahashi et al., 2013). To assess whether silencing PTSN could affect Rap1 phosphorylation, Rap1 was immunoprecipitated from cells in which endogenous PTSN expression was silenced by PTSN-specific shRNA. Rap1 phosphorylation was assessed by an antibody specific for phosphorylated PKA substrates. Silencing endogenous PTSN led to increased Rap1 phosphorylation and overexpression of shRNA resistant PTSN abolished Rap1 phosphorylation (Fig. 5A). To examine the specificity of increased phosphorylation by PTSN silencing, we examined cAMP Response Element-Binding Protein (CREB), an abundant PKA substrate. Silencing PTSN expression had no effect on Forskolin-induced CREB phosphorylation (Fig. 5B). Rap1a is phosphorylated at COOH-terminal Serine residue 180 by PKA both in vivo and in vitro(Takahashi et al., 2013). To directly address if Rap1 de-phosphorylation is important for RIAM MIT complex assembly, we asked if phosphorylation resistant Rap1a mutant (S180A) could support RIAM MIT assembly when PTSN was silenced. We generated IRES (internal ribosome entry site)-based bicistronic construct (Fig. 5C) expressing both Flag-RIAM and Myc-Rap1a(S180A) and transfected it into the cells expressing integrin α_IIb_β_3_ and Flag-RIAM. Expression of Rap1a(S180A) but not wild type Rap1a restored the association of integrin β_3_ with RIAM, indicative of the intact RIAM MIT complex, in PTSN silenced cells (Fig. 5C). Importantly, silencing PTSN did not affect the similar expression levels of Rap1a(S180A) or of Rap1a, nor did it affect expression of Flag-RIAM or α-tubulin (Fig. 5C). Together, these data show that PTSN stabilizes the MIT complex and thus, integrin activation by antagonizing the phosphorylation of Rap1.

**Figure 5.**
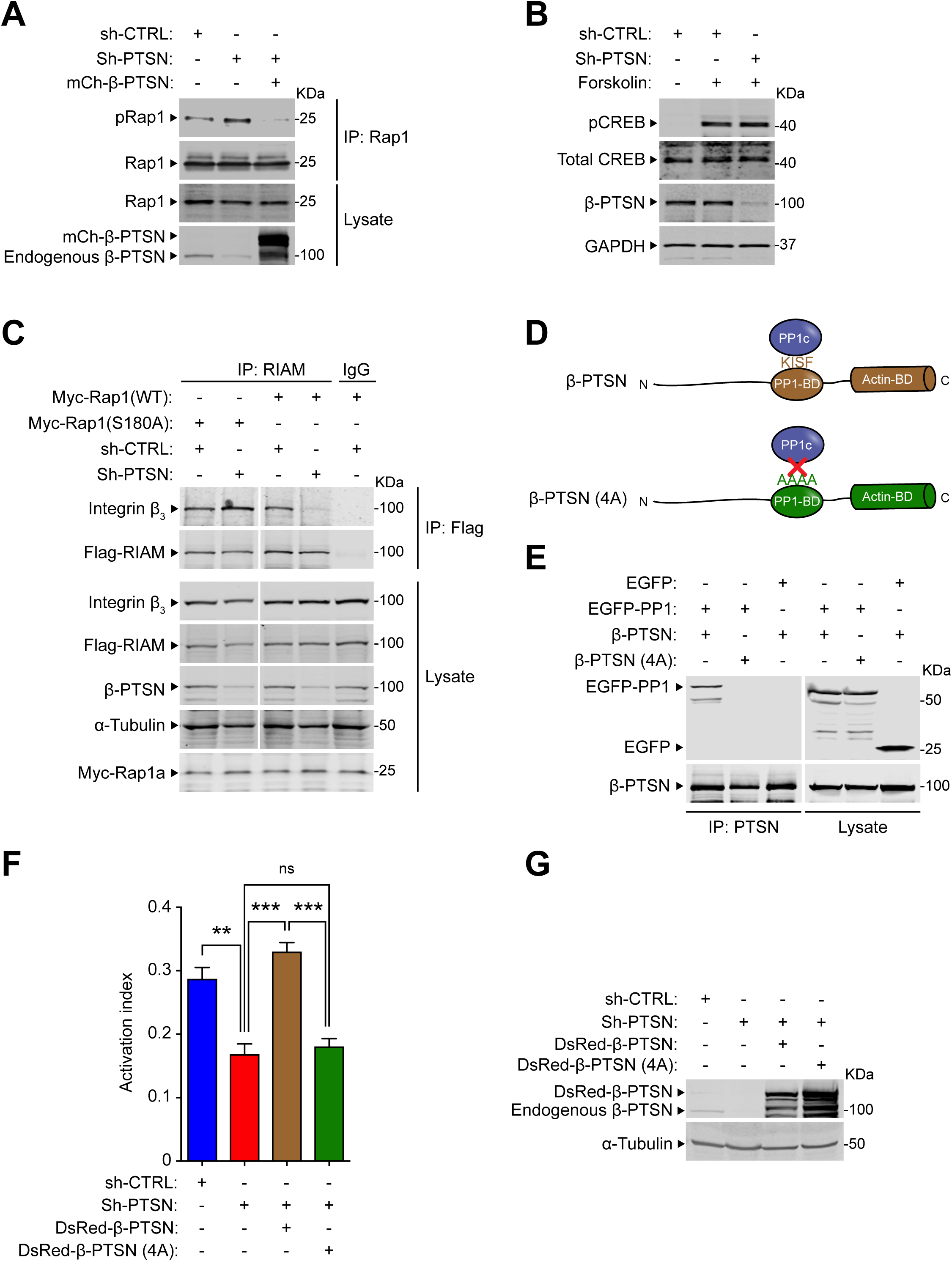
PTSN mediates Rap1 dephosphorylation. **(A)** Silencing of PTSN expression promotes Rap1 phosphorylation. α_IIb_(SBP)β_3_(D119A) expressing U2-OS cells were transduced with a lentivirus encoding a shRNA against PTSN or sh-CTRL (scrambled) and then transfected with a plasmid encoding Myc-tagged Rap1 in combination with a vector encoding mCherry-tagged PTSNor empty vector. Rap1 was immunoprecipitated using an anti-Myc antibody and phosphorylation of Rap1 was revealed by an antibody recognizing phosphorylated PKA substrates (pRap1). **(B)** As a control, the phosphorylation of CREB was assayed in PTSN-depleted cells using an antibody that specifically recognizes the phosphorylated form of CREB. U2-OS cells were treated with 10 μM forskolin for 30 min to induce CREB phosphorylation. **(C)** Phosphorylation-resistant Rap1 mutant (S180A) rescues RIAM MIT complex assembly. α_IIb_(SBP)β_3_(D119A) expressing U2-OS cells were transduced with a lentivirus encoding a shRNA against PTSN and transfected with a bicistronic plasmid encoding Myc-tagged Rap1 in combination with Flag-RIAM. RIAM was immunoprecipitated using an anti-Flag antibody and the associated β_3_ integrin was revealed by immunoblotting. **(D)** The KISF motif in PTSN binds to the catalytic subunit of protein phosphatase 1 (PP1c). **(E)** Mutation of the KISF residues into four Ala (4A) in PTSN blocks its interaction with PP1c. HEK293 cells were transfected with plasmids encoding EGFP-tagged PP1c in combination with either β-PTSN WT or (4A) mutant. PTSN was immunoprecipitated using an anti-Flag antibody prior to immunoblotting. **(F-G)** HEK293 cells that express constitutively active α_IIb_(R995A)β_3_ were transduced with a lentivirus encoding a shRNA against PTSN and transfected with a plasmid encoding EGFP-tagged shRNA resistant PTSN either WT or 4A mutant. **(F)** 293A cells that express constitutively active α_IIb_(R995A)β_3_ were transduced with a lentivirus encoding a shRNA against PTSN. A scrambled shRNA was used as a negative control. Cells were then transfected with a plasmid encoding DsRed-tagged shRNA resistant β-PTSN either WT or 4A mutant. Integrin activation was measured by flow cytometry using the monoclonal antibody PAC1 that specifically recognizes the activated form of α_IIb_β_3_. The activation index was calculated as (*F* - *Fo*)/(*Fm* - *Fo*), in which *F* is the mean fluorescence intensity (MFI) of PAC1 binding; *Fo* in presence of the competitive inhibitor integrilin; and *Fm* upon addition of the integrin-activating anti-LIBS6 antibody. Bar graphs represent mean ± SEM (n = 3 independent experiments). One-way ANOVA with Tukey post-test; ns not significant, ** p<0.01, *** p<0.001. **(G)** The expression of PTSN was confirmed by immunoblotting.

PTSN is a PP1 regulatory subunit and we confirmed that mutation of the PP1 catalytic subunit (PP1C) binding site disrupted its association with PTSN (Fig. 5E). This PP1C binding-defective mutant PTSN was unable to support integrin activation (Fig. 5F,G). PTSN is an actin associated protein(Lai et al., 2009; Wang et al., 2012), therefore we tested if silencing PTSN expression could affect the phosphorylation of regulators of actin dynamics whose activities are controlled by phosphorylation. Silencing of PTSN increased the phosphorylation of Cofilin (Supplementary Fig. 1A) and VASP (Supplementary Fig. 1B). Overexpression of shRNA resistant PTSN, either α or β isoforms, abolished the increased phosphorylation of cofilin (Supplementary Fig. 1C). Thus, in addition to preserving the activity of Rap1 and thus integrins, PTSN can induce de-phosphorylation of regulators of actin dynamics.

### *Ppp1r18^-/-^* mice are viable, fertile, and manifest reduced integrin activation

The foregoing data show that PTSN, a component of the MIT complex, stabilizes the assembly of that complex and resulting integrin activation by regulating the phosphorylation of Rap1. We therefore created *Ppp1r18^-/-^ mice* to assess a potential role for PTSN *in vivo.* We devised three different CRISPR/Cas9 strategies to inactivate *Ppp1r18*. We first used a combination of two different gRNAs to introduce 10 stop codons downstream of the codon for the initiator Methionine residue 428 of the α form of PTSN (Fig. 6A). To eliminate possible off target effects, we used different guide RNAs to delete the whole coding sequence of Exon 1 that contains initiator Methionine of both α and β forms of PTSN (Supplementary Fig. 2A). In both cases, homozygous *Ppp1r18^-/-^* offspring mice were viable and fertile and their splenocytes lacked both PTSN α and β isoforms (Fig. 6B, Supplementary Fig. 2B), and exhibited no change in expression of integrins α_L_, α_4_, β_2_, β_1_, or β_7_ (Fig. 6C) or of RIAM (Fig. 6D) in comparison with wild type littermates. To specifically inactivate the β form of PTSN, we used a gRNA upstream of the alternative translation start site at Methionine residue 428 resulting in a 10-base pair deletion which provokes a premature stop codon (Supplementary Fig. 3A) and absence of the β-PTSN form, but intact expression of the α form, in splenocytes (Supplementary Fig. 3B). Homozygotes of *Ppp1r18β^-/-^* mice, lacking only the β form of PTSN, were viable and fertile and their splenocytes expressed a normal complement of integrins α_L_, α_4_, β_2_, β_1_, or β_7_ (Supplementary Fig. 3C) or of RIAM (Supplementary Fig. 3D).

**Figure 6.**
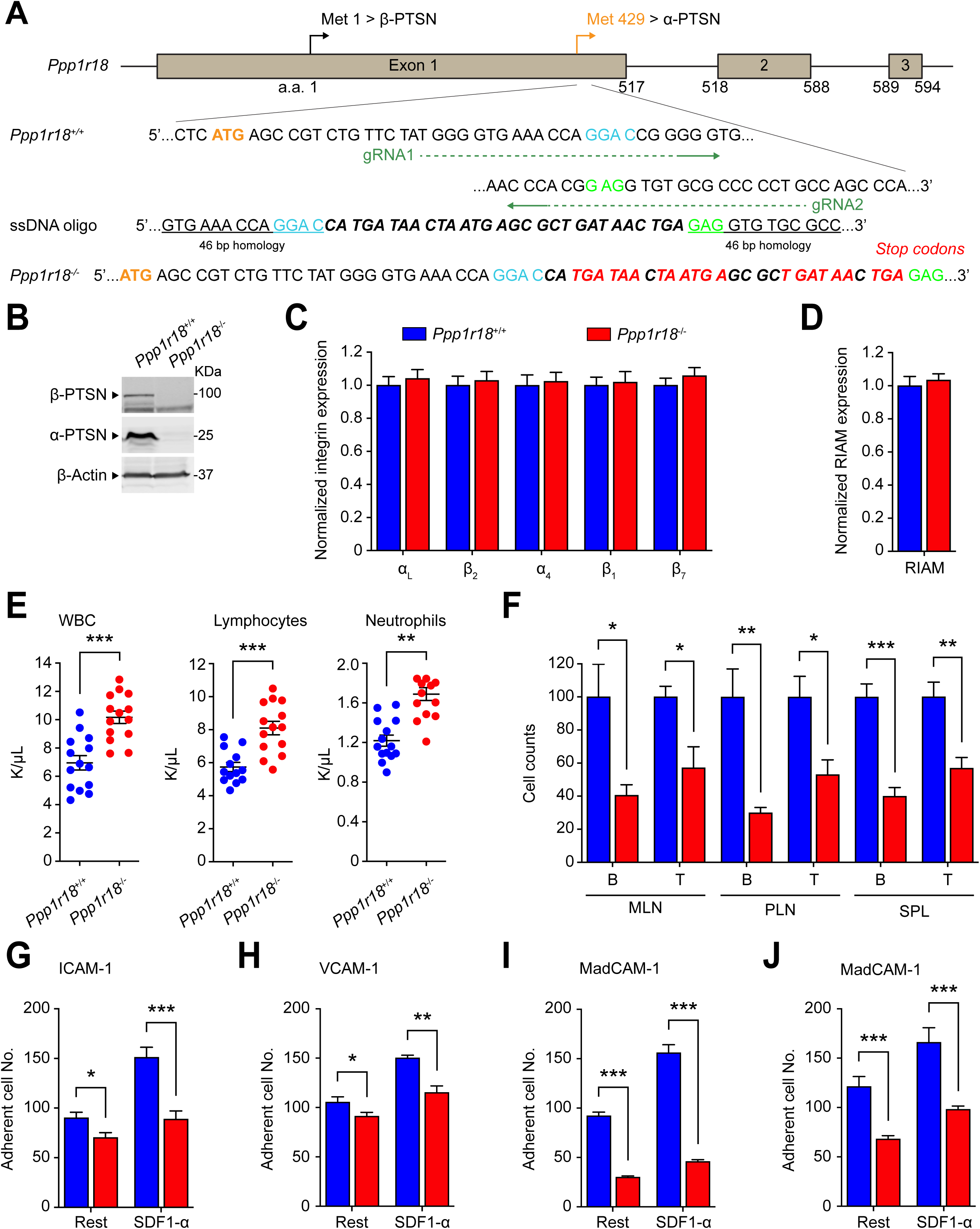
*Ppp1r18^-/-^* mice are viable with impaired T cell integrin activation. **(A)** Generation of *Ppp1r18^-/-^* mice. Two sgRNAs targeting the exon1 downstream Methionine residue 429, which drives the expression of α-PTSN, were used in combination with a ssDNA oligo to introduce seven stop codons. **(B)** *Ppp1r18^-/-^* mice manifest loss of both α- and β-PTSN isoforms. PTSN expression in splenocytes was assayed by immunoblotting using an antibody that reacts with the C-terminus of PTSN and recognizes both α- and β-PTSN isoforms. **(C)** Surface expression of α_L_ (CD11a), β_2_ (CD18), α_4_ (CD49d), β_1_ (CD29) and β_7_ integrins, and **(D)** intracellular staining of RIAM in splenocytes was measured by flow cytometry. Bar graphs represent mean ± SEM (n = 5 mice). Data are normalized to *Ppp1r18^+/+^* samples. Two-tailed *t*-test; no significant differences were observed. **(E-F)** *Ppp1r18^-/-^* mice exhibit a leukocytosis. **(E)** Peripheral blood cell counts of WT or *Ppp1r18^-/-^* mice. Mean ± SEM are plotted. Two-tailed *t*-test; ** p<0.01; *** p<0.001. **(F)** The number of T and B cells in mesenteric lymph node (MLN), peripheral lymph node (PLN) and spleen (SPL) from WT or *Ppp1r18^-/-^* mice. Data are normalized to *Ppp1r18^+/+^* samples. Bar graphs represent mean ± SEM (n = 4 mice). Two-tailed *t*-test; * p<0.05, ** p<0.01, *** p<0.001. **(G-J)** Loss of PTSN reduces T and B cell adhesion. CD4^+^ T cells **(G-I)** or B cells **(J)** were isolated from the spleen of WT or *Ppp1r18^-/-^* mice. Cell adhesion to immobilized ICAM-1, VCAM-1 or MAdCAM-1 was assayed in flow condition upon stimulation with SDF-1α. Rest, no stimulation. Bar graphs represent mean ± SEM (n = 5 mice). Two-tailed *t*-test; * p<0.05, ** p<0.01, *** p<0.001.

*Ppp1r18^-/-^* mice of both genotypes lacking both forms of PTSN, exhibited a ∼two-fold increase in blood leukocytes that affected both lymphocytes and neutrophils (Fig. 6E) and an accompanying decrease in T and B cells in peripheral lymphatic organs (Fig. 6F). The leukocytosis in these mice combined with a reduction in lymphocytes in peripheral lymphatic organs is similar to that observed in RIAM deficient *Apbb1ip^-/-^* mice(Klapproth et al., 2015) (Su et al., 2015) and is explained by the reduction in T cell (Fig. 6G-I) and B cell (Fig. 6J) integrin activation. Furthermore, *Ppp1r18^-/-^* splenocytes exhibited increased phosphorylation of cofilin (Supplementary Fig. 1D) and VASP (Supplementary Fig. 1E) similar to that observed in U2-OS cells (Supplementary Fig. 1A,B). We were unable to immunoprecipitate sufficient Rap1 from splenocytes to analyze its phosphorylation. Thus, PTSN enables activation of lymphocyte integrins and population of peripheral lymphoid organs.

In sharp contrast to *Ppp1r18^-/-^* mice, *Ppp1r18β^-/-^* mice exhibited blood counts indistinguishable from those of wild type littermates (Supplementary Fig. 3E). Consistent with the absence of leukocytosis, *Ppp1r18β^-/-^* T cells exhibited similar activation of T cell α_L_β_2_ and α_4_β_1_ integrins as judged by adhesion on ICAM-1, or VCAM-1 respectively (Supplementary Fig. 3G,H). The preservation of lymphocyte integrin function was manifested phenotypically by the normal population of peripheral lymphatic tissues by B cells and T cells (Supplementary Fig. 3F). Because *Ppp1r18β^-/-^* mice exhibited no obvious phenotype, α-PTSN is sufficient for these lymphocyte functions.

### *Ppp1r18^-/-^* T cells exhibit reduced capacity to induce colitis

Because PTSN null mice were apparently healthy, yet exhibited defects in T cell integrin activation, we hypothesized that their T cells might lack the capacity to provoke auto-immune inflammation. We induced experimental autoimmune colitis(Sun et al., 2020) by infusing CD4^+^CD25^-^CD45RB^high^ T cells (T_CONV_) from *Ppp1r18^-/-^* mice or WT mice into *Rag1^-/-^* recipient mice. *Rag1^-/-^* mice injected with WT T_CONV_ manifested a progressive loss in bodyweight 20-30 days after the infusion (Fig. 7A). Half of these mice died by 100 days (Fig. 7B). In contrast, *Rag1^-/-^* mice injected with *Ppp1r18^-/-^* T_CONV_ exhibited a little loss of bodyweight and only 10 % of these mice died by 100 days (Fig. 7A,B).

**Figure 7.**
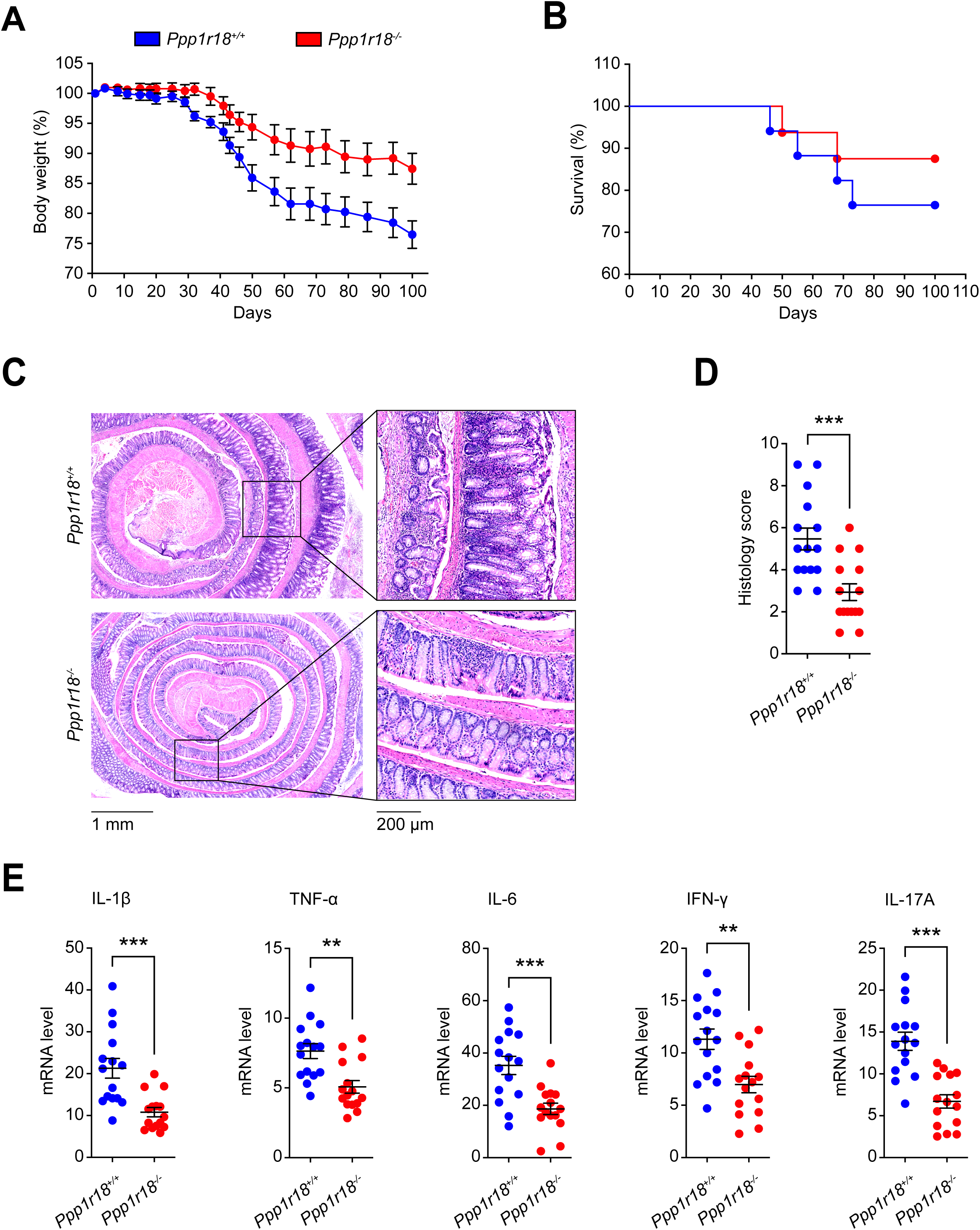
Loss of PTSN expression protects mice from colitis. Conventional T cells (1.10^6^) from WT or *Ppp1r18^-/-^* mice were adoptively transferred into *Rag1^-/-^* mice. **(A)** Changes in body weight are shown. Values are normalized as a percentage of the original weight. **(B)** Percent survival. Significant differences were determined using a two-way ANOVA with Bonferroni post-test. **(C)** Representative hematoxylin and eosin staining of distal colon sections wrapped as swiss rolls from *Rag1^-/-^* mice injected with conventional T cells from WT or *Ppp1r18^-/-^* mice at Day 100. Scale bars: 1 mm or 200 μm. **(D)** Histology score. Two-tailed *t*-test; *** p<0.001. **(E)** mRNA expression of IL-1β, TNF-α, IL-6, IFN-γ and IL-17A in distal colon tissues at Day 100. Results are normalized to expression of GAPDH. Mean ± SEM are plotted. Two-tailed *t*-test; ** p<0.01, *** p<0.001.

Histologically, wild-type T_CONV_ infusion led to a severe colitis in *Rag1^-/-^* mice, with almost complete loss of crypts, dense infiltrates of leukocytes in both mucosa and submucosa, and thickening of the bowel wall (Fig. 7C). By contrast, the inflammatory infiltrates in the *Rag1^-/-^* mice infused with *Ppp1r18^-/-^* T_CONV_ were much reduced and less tissue damage was observed (Fig. 7C). Blinded histological scoring for inflammatory cell infiltrates and epithelial damage confirmed a reduction in the severity of colitis in *Ppp1r18^-/-^* T_CONV_ infused *Rag1^-/-^* mice (Fig. 7D). The difference in inflammatory cell infiltration between *Ppp1r18^-/-^* and wild-type T_CONV_ infused *Rag1^-/-^* mice was confirmed by the reduction in colonic expression of pro-inflammatory cytokines (IL-1β, TNF-α, IL-6, IFN-γ and IL-17A) in *Ppp1r18^-/-^* T_CONV_ recipients (Fig. 7E). Thus, the defect in integrin activation in PTSN null CD4^+^ T cells is associated with impaired capacity to induce intestinal inflammation.

## DISCUSSION

The MIT complex plays an essential role in leukocyte integrin activation and thus in the trafficking and functions of lymphocytes in immunity and immunopathology(Su et al., 2015). Here we report that an actin binding regulatory subunit of PP1, PTSN, encoded by the *Ppp1r18* gene(Kao et al., 2007), stabilizes the MIT complex by mediating the dephosphorylation of Rap1 thereby preserving Rap1 activity. The stabilization of the MIT complex enables lymphocyte integrin activation and cell adhesion. *Ppp1r18^-/-^* mice are viable, fertile, and apparently healthy. These mice exhibit reduced lymphocyte population of peripheral lymphoid tissues associated with lymphocytosis in peripheral blood, findings ascribable to defective activation of lymphocyte integrins α_L_β_2_ and α_4_β_7_. Furthermore, *Ppp1r18^-/-^* T cells manifested reduced capacity to induce colitis in an adoptive transfer model. Thus, PTSN is a regulator of T cell integrin-mediated functions that stabilizes the RIAM MIT complex by dephosphorylating Rap1. In spite of this defect in T cell integrin function *Ppp1r18^-/-^* mice appear healthy suggesting that PTSN may be a therapeutic target in T cell mediated autoimmune diseases.

PTSN is physically associated with MIT complex and enables integrin activation by mediating de-phosphorylation of Rap1, thereby stabilizing the complex. PTSN was associated with the MIT complex formed by a ligand binding-defective integrin(Loftus et al., 1990) explaining why this association can form in non-adherent lymphocytes. Activation of integrins in lymphocytes requires Rap1(Su et al., 2015) and we find that PTSN helps maintain Rap1 in a functional non-phosphorylated state. In particular, we find that silencing of phostensin destabilizes the MIT complex and this destabilization can be prevented by a non-phosporylatable mutant of Rap1a. In addition to blocking membrane association of Rap1, Rap1 phosphorylation results in prolonged activation of ERK 1/2(Takahashi et al., 2017), an event that can also suppress integrin activation(Hughes et al., 1997) and assembly of the MIT complex(Lagarrigue et al., 2015). Thus, PTSN joins Phactr4(Zhang et al., 2012) as actin-associated protein phosphatase 1 regulators that modulate integrin functions.

*Ppp1r18^-/-^* mice exhibited lymphocytosis accompanied by a reduced population of peripheral lymphoid organs, a phenotype indicative of a failure of lymphocyte migration out of the blood as a consequence of reduced integrin activation. Cell migration requires anterior-posterior polarization of processes such as integrin activation and of actin dynamics to support directional migration(Ridley et al., 2003). The MIT complex is localized to and mediates the protrusions that form at the leading edge of migrating cells(Lagarrigue et al., 2015; Lee et al., 2013) , a site at which α4 integrins can anchor Type I Protein Kinase A (PKA)(Lim et al., 2007; Lim et al., 2008) to enable PKA activation. PKA phosphorylation of RhoA forms protrusion-retraction pacemaker at the leading edge of migrating cells(Tkachenko et al., 2011). PTSN, by opposing PKA-mediated Rap1 phosphorylation, can preserve stability of the MIT complex at sites of increased PKA activity such as the leading edge(Lim et al., 2008). Conversely, over-expression of phospho-resistant Rap1 can arrest migration(Takahashi et al., 2013), suggesting that the dynamic interplay between PKA and PTSN in Rap1 phosphorylation can contribute to the protrusion-retraction cycles that govern cell migration.

Pre-clinical research identified inhibition of ligand binding to lymphocyte integrins as a potential therapeutic target in autoimmune and inflammatory diseases(Dustin, 2019; von Andrian and Engelhardt, 2003). This idea was validated by the success of vedolizumab anti-α_4_β_7_ in IBD(Feagan et al., 2013), natalizumab anti-α_4_β_1_ in multiple sclerosis(Rice et al., 2005), and efalizumab anti-α_L_β_2_ in psoriasis(Dedrick et al., 2002). Serious mechanisms-based toxicities such as progressive multifocal leukoencephalopathy have limited the use of natalizumab and efalizumab. Inhibition of integrin signaling can preserve some integrin function and can therefore ameliorate mechanism-based toxicities of complete blockade of integrin function(Feral et al., 2006; Petrich et al., 2007). Talin and Rap1 are major elements in signaling pathways that activate leukocyte integrins; however, global loss of talin-1 or combined loss of Rap1a and Rap1b, leads to embryonic lethality in mice(Calderwood et al., 2013; Li et al., 2007). RIAM plays a key role in Rap1- dependent talin-mediated activation of α_L_β_2_ and α_4_β_7_ integrins(Su et al., 2015; Sun et al., 2021) in most leukocytes and lack of RIAM leads to no obvious developmental defects or abnormalities in platelet functions(Klapproth et al., 2015; Stritt et al., 2015; Su et al., 2015). Here we show that PTSN supports Rap1- induced assembly of the RIAM-talin-integrin complex required for integrin activation. In the absence of PTSN a defect in lymphocyte trafficking is associated with the inability of T cells to provoke experimental colitis. Importantly, like RIAM, PTSN is expressed at low levels in platelets (Fig. 3A) and we observed no gross bleeding tendency or anemia in *Apbb1ip^-/-^* mice. Thus, PTSN inhibition may provide a means to blunt immune-mediated inflammation with less mechanism-based toxicity than direct blockade of ligand binding to leukocyte integrins.

## Acknowledgements

We thank the UCI and the UCSD Transgenic Mouse Facilities for design help and production of CRISPR modified mice; Jennifer Santini for microscopy technical assistance; and the UCSD School of Medicine Microscopy Core (NINDS P30NS047101). The UCI TMF is a shared resource funded in part by the Chao Family Comprehensive Cancer Center Support Grant (P30CA062203) from the National Cancer Institute. This work was supported by grants HL 139947 and HL 151433 to MHG and by the American Heart Association Career Development Award 18CDA34110228 (F.L.) and Scientist Development Grant 14SDG18440023 (HSL).

## Author Contributions

M.H.G conceived the study, designed experiments, analyzed data, and wrote the manuscript. H.-S.L., H.S. and F.L. designed and executed experiments, analyzed data, and wrote the manuscript. J.W.F and N.E.S. performed mass spectroscopic analysis. A.R.G designed and produced antigen, and tested the antibody produced. M.H.G, H.-S.L., H.S., F.L., and A.R.G edited the manuscript.

## SUPPLEMENTARY FIGURE LEGENDS

**Figure S1.**
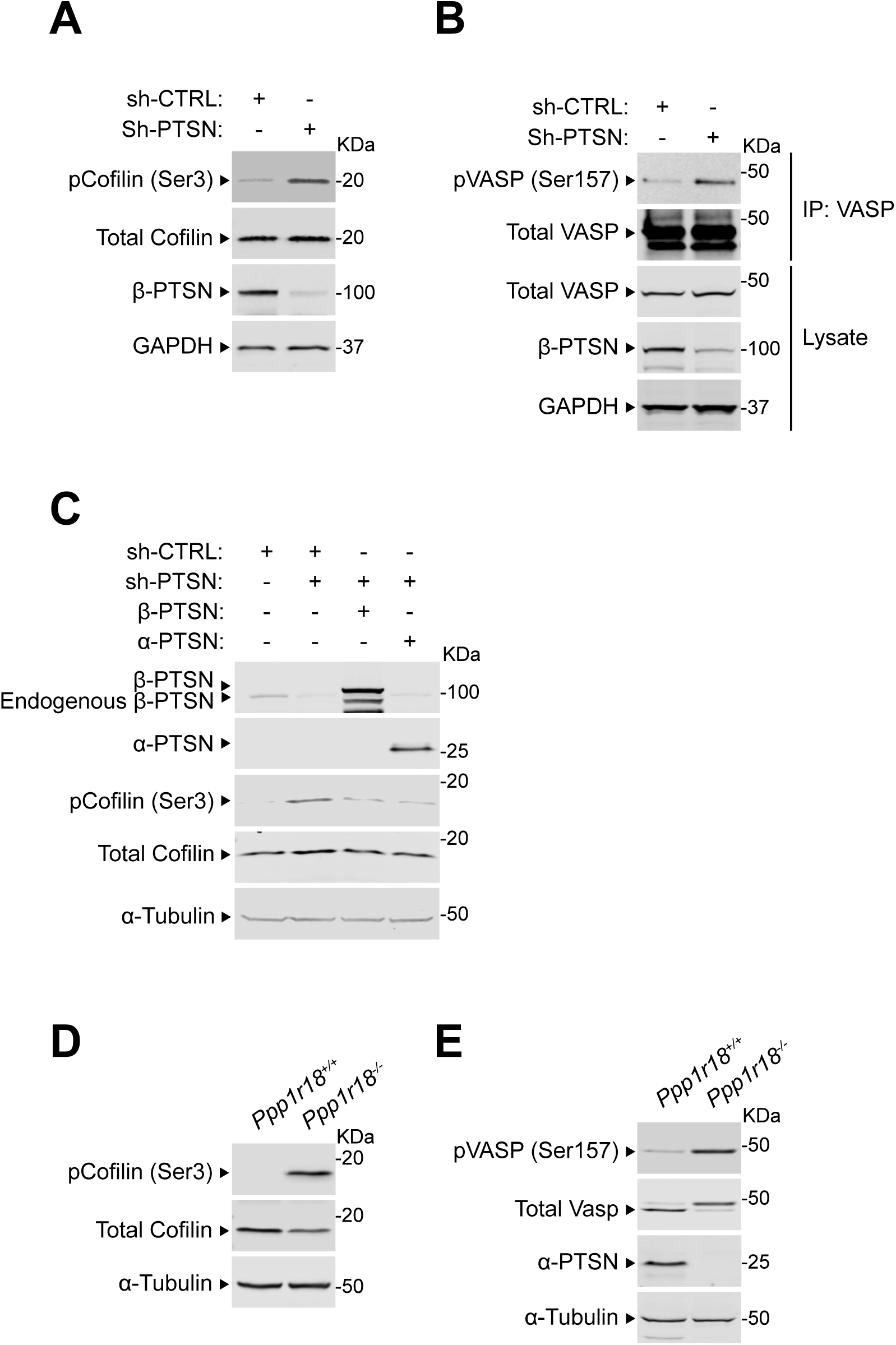
PTSN regulates the phosphorylation of Cofilin and VASP. **(A-C)** U2-OS cells were transduced with a lentivirus encoding a shRNA against PTSN. A scrambled shRNA was used as a negative control. **(B)** Endogenous VASP was captured by immunoprecipitation. **(C)** Cells were transfected with a plasmid encoding either Flag-tagged α- or β-PTSN isoforms. **(D-E)** Lysates of splenocytes isolated from WT or *Ppp1r18^-/-^* mice. **(A-E)** Phosphorylation of Cofilin and VASP were analyzed by immunoblotting using antibodies that specifically recognize the phosphoproteins. Results are representative of 3 independent experiments.

**Figure S2.**
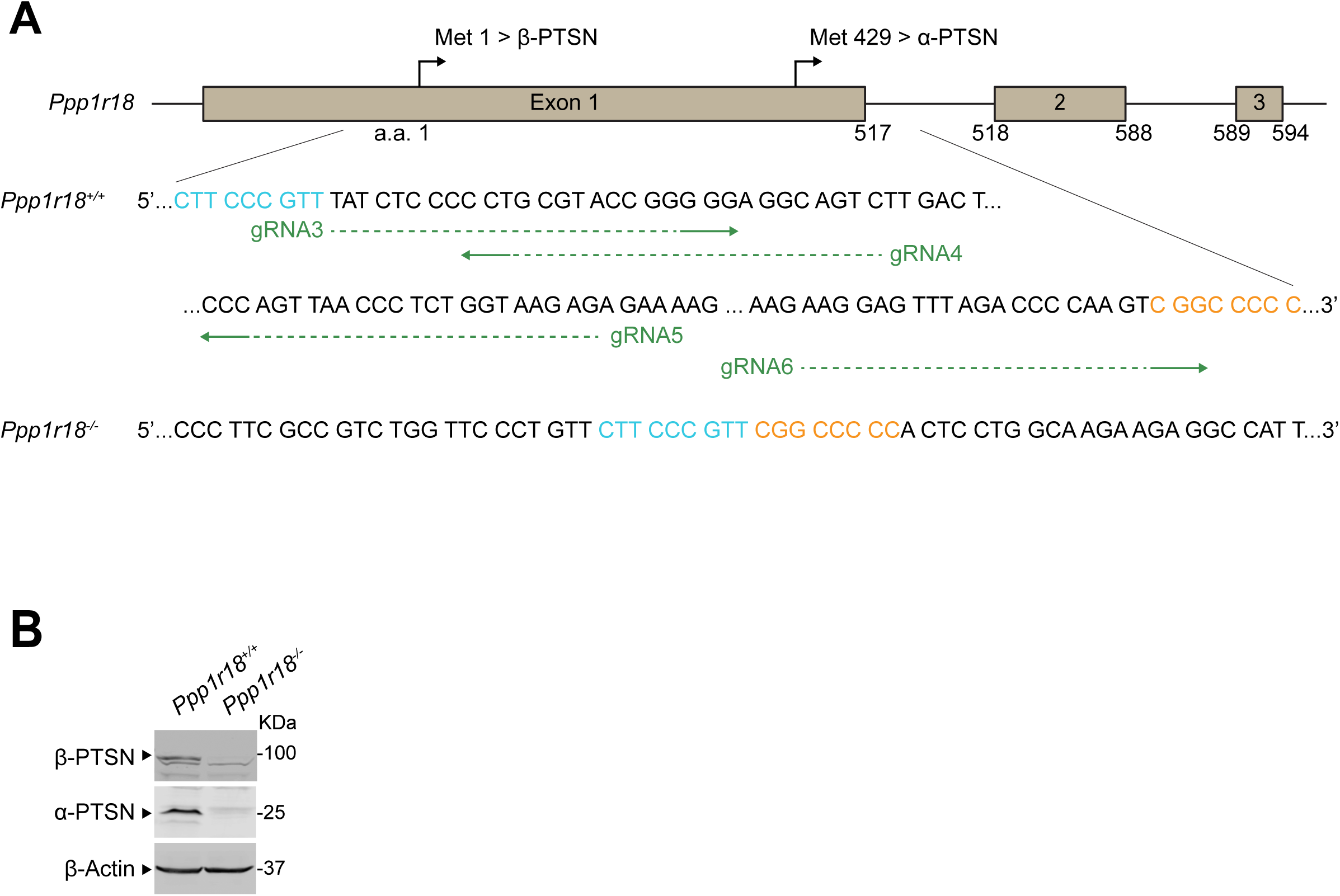
Alternative generation of *Ppp1r18^-/-^* mice. **(A)** Generation of *Ppp1r18^-/-^* mice by deleting the whole coding sequence of exon1. Four sgRNAs targeting the exon1 were used. The edited sequence of the repaired *Ppp1r18^-/-^* allele is shown. **(B)** *Ppp1r18^-/-^* mice manifest loss of both α- and β-PTSN isoforms. PTSN expression in splenocytes was assayed by immunoblotting using an antibody that reacts with the C-terminus of PTSN and recognizes both α- and β- PTSN isoforms.

**Figure S3.**
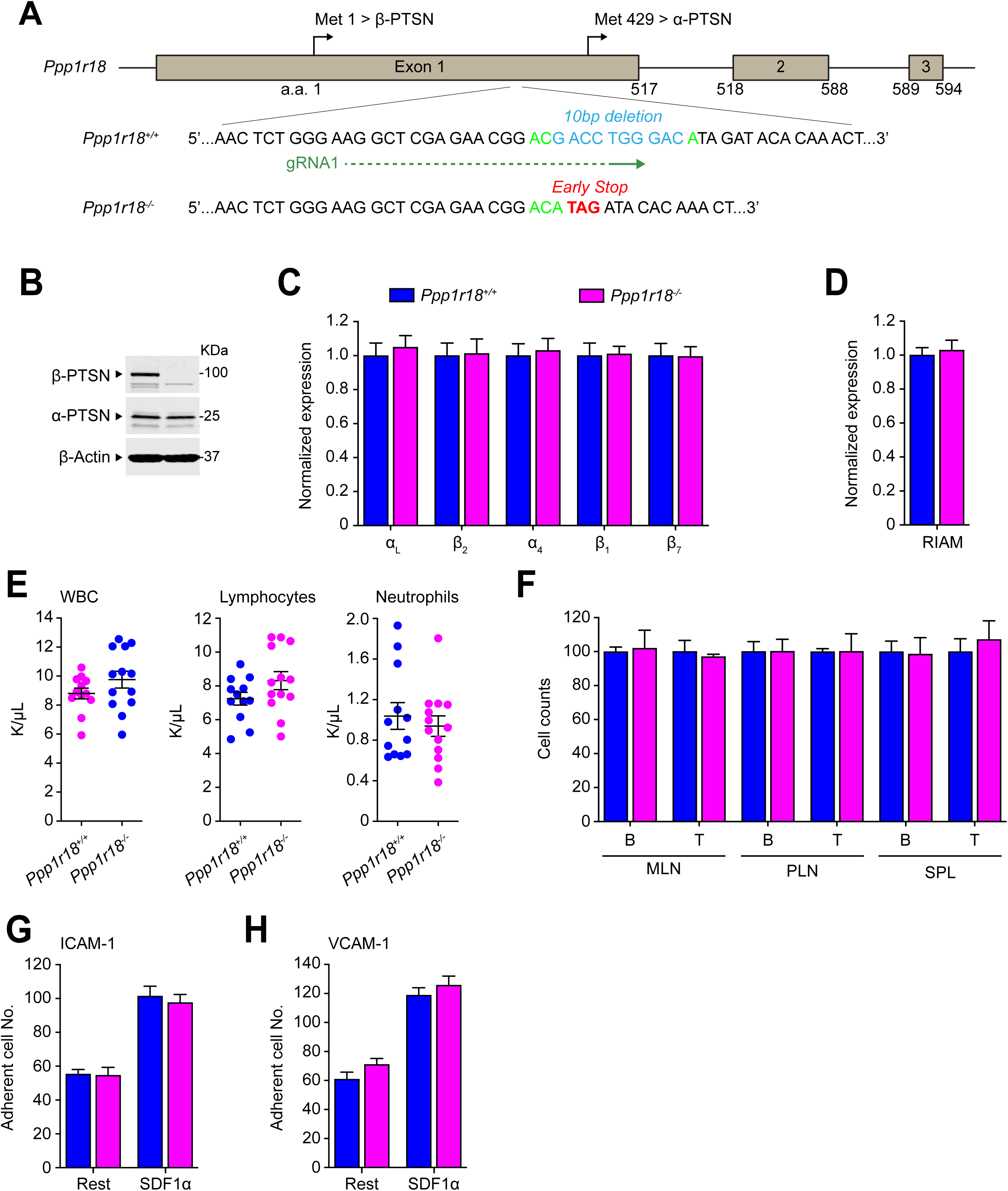
Deletion of the PTSN β isoform in mice leads to intact T cell integrin activation. **(A)** Generation of *Ppp1r18β^-/-^* mice. Two sgRNAs targeting the exon1 in between Methionine residues 1 and 429, which drive the expression of β- and α-PTSN isoforms, respectively. Edition led to a 10-base pair deletion resulting in an early stop codon. **(B)** *Ppp1r18β^-/-^* mice manifest loss of the PTSN β isoform, but not the α-PTSN isoform. PTSN expression in splenocytes was assayed by immunoblotting using an antibody that reacts with the C-terminus of PTSN and recognizes both α- and β-PTSN isoforms. **(C-D)** Surface expression of α_L_ (CD11a), β_2_ (CD18), α_4_ (CD49d), β_1_ (CD29) and β_7_ integrins **(C)** or intracellular staining of RIAM **(D)** in splenocytes was measured by flow cytometry. Bar graphs represent mean ± SEM (n = 3 mice). Data are normalized to *Ppp1r18β^+/+^* samples. Two-tailed *t*-test; no significant differences were observed. **(E)** *Ppp1r18β^-/-^* mice exhibit normal peripheral blood cell counts. Mean ± SEM are plotted. Two-tailed *t*-test; no significant differences were observed. **(F)** The number of T and B cells in mesenteric lymph node (MLN), peripheral lymph node (PLN) and spleen (SPL) from WT or *Ppp1r18β^-/-^* mice. Data are normalized to *Ppp1r18β^+/+^* samples. Bar graphs represent mean ± SEM (n = 4 mice). Two-tailed *t*-test; no significant differences were observed. **(G-H)** Deletion of *Ppp1r18β^-/-^* does not impair T cell adhesion. CD4^+^ T cells were isolated from the spleen of WT or *Ppp1r18β^-/-^* mice. Cell adhesion to immobilized ICAM-1 or VCAM-1 was assayed in flow condition upon stimulation with SDF-1α. Rest, no stimulation. Bar graphs represent mean ± SEM (n = 6 mice). Two-tailed *t*-test; no significant differences were observed.

## REFERENCES

Bos, J.L. (2005). Linking Rap to cell adhesion. CurrOpinCell Biol 17, 123.

Boussiotis, V.A., Freeman, G.J., Berezovskaya, A., Barber, D.L., and Nadler, L.M. (1997). Maintenance of human T cell anergy: blocking of IL-2 gene transcription by activated Rap1. Science 278, 124.

Calderwood, D.A., Campbell, I.D., and Critchley, D.R. (2013). Talins and kindlins: partners in integrin-mediated adhesion. Nature reviews Molecular cell biology 14, 503–517.

Dedrick, R.L., Walicke, P., and Garovoy, M. (2002). Anti-adhesion antibodies efalizumab, a humanized anti-CD11a monoclonal antibody. Transpl Immunol 9, 181–186.

Dustin, M.L. (2019). Integrins and Their Role in Immune Cell Adhesion. Cell 177, 499–501.

Erben, U., Loddenkemper, C., Doerfel, K., Spieckermann, S., Haller, D., Heimesaat, M.M., Zeitz, M., Siegmund, B., and Kuhl, A.A. (2014). A guide to histomorphological evaluation of intestinal inflammation in mouse models. Int J Clin Exp Pathol 7, 4557–4576.

Feagan, B.G., Rutgeerts, P., Sands, B.E., Hanauer, S., Colombel, J.F., Sandborn, W.J., Van Assche, G., Axler, J., Kim, H.J., Danese, S., et al. (2013). Vedolizumab as induction and maintenance therapy for ulcerative colitis. N Engl J Med 369, 699–710.

Feral, C.C., Rose, D.M., Han, J., Fox, N., Silverman, G.J., Kaushansky, K., and Ginsberg, M.H. (2006). Blocking the alpha 4 integrin-paxillin interaction selectively impairs mononuclear leukocyte recruitment to an inflammatory site. J ClinInvest 116, 715.

Frelinger, A.L., III, Cohen, I., Plow, E.F., Smith, M.A., Roberts, J., Lam, S.C.T., and Ginsberg, M.H. (1990). Selective inhibition of integrin function by antibodies specific for ligand-occupied receptor conformers. Journal of Biological Chemistry 265, 6346.

Frelinger, A.L., III, Du, X., Plow, E.F., and Ginsberg, M.H. (1991). Monoclonal antibodies to ligand-occupied conformers of integrin alpha IIb beta 3 (Glycoprotein IIb-IIIa) alter receptor affinity, specificity, and function. Journal of Biological Chemistry 266, 17106.

Frelinger, A.L., III, Lam, S.C.T., Plow, E.F., Smith, M.A., Loftus, J.C., and Ginsberg, M.H. (1988). Occupancy of an adhesive glycoprotein receptor modulates expression of an antigenic site involved in cell adhesion. Journal of Biological Chemistry 263, 12397.

Goldfinger, L.E., Ptak, C., Jeffery, E.D., Shabanowitz, J., Han, J., Haling, J.R., Sherman, N.E., Fox, J.W., Hunt, D.F., and Ginsberg, M.H. (2007). An Experimentally Derived Database of Candidate Ras-Interacting Proteins. J ProteomeRes 6, 1806.

Hogg, N., Patzak, I., and Willenbrock, F. (2011). The insider’s guide to leukocyte integrin signalling and function. Nature reviews Immunology 11, 416–426.

Hughes, P.E., Renshaw, M.W., Pfaff, M., Forsyth, J., Keivens, V.M., Schwartz, M.A., and Ginsberg, M.H. (1997). Suppression of integrin activation: A novel function of a Ras/Raf- initiated MAP kinase pathway. Cell 88, 521.

Hynes, R.O. (2002). Integrins: bidirectional, allosteric signaling machines. Cell 110, 673–687.

Kao, S.C., Chen, C.Y., Wang, S.L., Yang, J.J., Hung, W.C., Chen, Y.C., Lai, N.S., Liu, H.T., Huang, H.L., Chen, H.C., et al. (2007). Identification of phostensin, a PP1 F-actin cytoskeleton targeting subunit. Biochemical and biophysical research communications 356, 594–598.

Kim, C., Ye, F., and Ginsberg, M.H. (2011). Regulation of integrin activation. Annual review of cell and developmental biology 27, 321–345.

Klapproth, S., Sperandio, M., Pinheiro, E.M., Prunster, M., Soehnlein, O., Gertler, F.B., Fassler, R., and Moser, M. (2015). Loss of the Rap-1 effector RIAM results in leukocyte adhesion deficiency due to impaired beta2 integrin function in mice. Blood 126, 2704–2712.

Lagarrigue, F., Gertler, F.B., Ginsberg, M.H., and Cantor, J.M. (2017). Cutting Edge: Loss of T Cell RIAM Precludes Conjugate Formation with APC and Prevents Immune-Mediated Diabetes. Journal of immunology (Baltimore, Md : 1950) 198, 3410–3415.

Lagarrigue, F., Vikas Anekal, P., Lee, H.S., Bachir, A.I., Ablack, J.N., Horwitz, A.F., and Ginsberg, M.H. (2015). A RIAM/lamellipodin-talin-integrin complex forms the tip of sticky fingers that guide cell migration. Nature communications 6, 8492.

Lai, N.S., Wang, T.F., Wang, S.L., Chen, C.Y., Yen, J.Y., Huang, H.L., Li, C., Huang, K.Y., Liu, S.Q., Lin, T.H., et al. (2009). Phostensin caps to the pointed end of actin filaments and modulates actin dynamics. Biochemical and biophysical research communications 387, 676–681.

Lee, H.S., Anekal, P., Lim, C.J., Liu, C.C., and Ginsberg, M.H. (2013). Two Modes of Integrin Activation Form a Binary Molecular Switch in Adhesion Maturation. Molecular biology of the cell 24, 1354–1362.

Lefort, C.T., Rossaint, J., Moser, M., Petrich, B.G., Zarbock, A., Monkley, S.J., Critchley, D.R., Ginsberg, M.H., Fassler, R., and Ley, K. (2012). Distinct roles for talin-1 and kindlin-3 in LFA-1 extension and affinity regulation. Blood 119, 4275–4282.

Ley, K., Laudanna, C., Cybulsky, M.I., and Nourshargh, S. (2007). Getting to the site of inflammation: the leukocyte adhesion cascade updated. Nature reviews Immunology 7, 678–689.

Ley, K., Rivera-Nieves, J., Sandborn, W.J., and Shattil, S. (2016). Integrin-based therapeutics: biological basis, clinical use and new drugs. Nature reviews Drug discovery 15, 173–183.

Li, Y., Yan, J., De, P., Chang, H.C., Yamauchi, A., Christopherson, K.W., 2nd, Paranavitana, N.C., Peng, X., Kim, C., Munugalavadla, V., et al. (2007). Rap1a null mice have altered myeloid cell functions suggesting distinct roles for the closely related Rap1a and 1b proteins. Journal of immunology (Baltimore, Md : 1950) 179, 8322–8331.

Lim, C.J., Han, J., Yousefi, N., Ma, Y., Amieux, P.S., McKnight, G.S., Taylor, S.S., and Ginsberg, M.H. (2007). alpha4 Integrins are Type I cAMP-dependent protein kinase-anchoring proteins. Nat Cell Biol 4, 415.

Lim, C.J., Kain, K.H., Tkachenko, E., Goldfinger, L.E., Gutierrez, E., Allen, M.D., Groisman, A., Zhang, J., and Ginsberg, M.H. (2008). Integrin-mediated protein kinase A activation at the leading edge of migrating cells. Molecular biology of the cell 19, 4930–4941.

Lin, Y.S., Huang, H.L., Liu, W.T., Lin, T.H., and Huang, H.B. (2014). Identification of the high molecular weight isoform of phostensin. International journal of molecular sciences 15, 1068–1079.

Lin, Y.S., Huang, K.Y., Wang, T.F., Huang, H.L., Yu, H.C., Yen, J.Y., Hung, S.H., Liu, S.Q., Lai, N.S., and Huang, H.B. (2011). Immunolocalization of phostensin in lymphatic cells and tissues. The journal of histochemistry and cytochemistry : official journal of the Histochemistry Society 59, 741–749.

Loftus, J.C., O’Toole, T.E., Plow, E.F., Glass, A., Frelinger, A.L., III, and Ginsberg, M.H. (1990). A β3 integrin mutation abolishes ligand binding and alters divalent cation-dependent conformation. Science 249, 915.

Luo, B.H., Carman, C.V., and Springer, T.A. (2007). Structural basis of integrin regulation and signaling. Annual review of immunology 25, 619–647.

Medrano-Fernandez, I., Reyes, R., Olazabal, I., Rodriguez, E., Sanchez-Madrid, F., Boussiotis, V.A., Reche, P.A., Cabanas, C., and Lafuente, E.M. (2013). RIAM (Rap1-interacting adaptor molecule) regulates complement-dependent phagocytosis. Cellular and molecular life sciences : CMLS 70, 2395–2410.

Mombaerts, P., Iacomini, J., Johnson, R.S., Herrup, K., Tonegawa, S., and Papaioannou, V.E. (1992). RAG-1-deficient mice have no mature B and T lymphocytes. Cell 68, 869–877.

O’Toole, T.E., Katagiri, Y., Faull, R.J., Peter, K., Tamura, R., Quaranta, V., Loftus, J.C., Shattil, S.J., and Ginsberg, M.H. (1994). Integrin cytoplasmic domains mediate inside-out signal transduction. The Journal of cell biology 124, 1047–1059.

Petrich, B.G., Fogelstrand, P., Partridge, A.W., Yousefi, N., Ablooglu, A.J., Shattil, S.J., and Ginsberg, M.H. (2007). The antithrombotic potential of selective blockade of talin-dependent integrin alphaIIbbeta3 (platelet GPIIb-IIIa) activation. Journal of Clinical Investigation 117, 2250–2259.

Rice, G.P., Hartung, H.P., and Calabresi, P.A. (2005). Anti-alpha4 integrin therapy for multiple sclerosis: mechanisms and rationale. Neurology 64, 1336–1342.

Ridley, A.J., Schwartz, M.A., Burridge, K., Firtel, R.A., Ginsberg, M.H., Borisy, G., Parsons, J.T., and Horwitz, A.R. (2003). Cell migration: integrating signals from front to back. Science 302, 1704.

Rose, D.M., Cardarelli, P.M., Cobb, R.R., and Ginsberg, M.H. (2000). Soluble VCAM-1 binding to alpha 4 integrins is cell type-specific, activation-dependent, and disrupted during apoptosis. Blood 95, 602.

Shattil, S.J., Hoxie, J.A., Cunningham, M., and Brass, L.F. (1985). Changes in the platelet membrane glycoprotein IIb-IIIa Complex during platelet activation. Journal of Biological Chemistry 260, 11107.

Stritt, S., Wolf, K., Lorenz, V., Vogtle, T., Gupta, S., Bosl, M.R., and Nieswandt, B. (2015). Rap1-GTP-interacting adaptor molecule (RIAM) is dispensable for platelet integrin activation and function in mice. Blood 125, 219–222.

Su, W., Wynne, J., Pinheiro, E.M., Strazza, M., Mor, A., Montenont, E., Berger, J., Paul, D.S., Bergmeier, W., Gertler, F.B., et al. (2015). Rap1 and its effector RIAM are required for lymphocyte trafficking. Blood 126, 2695–2703.

Sun, H., Kuk, W., Rivera-Nieves, J., Lopez-Ramirez, M.A., Eckmann, L., and Ginsberg, M.H. (2020). beta7 Integrin Inhibition Can Increase Intestinal Inflammation by Impairing Homing of CD25(hi)FoxP3(+) Regulatory T Cells. Cellular and molecular gastroenterology and hepatology 9, 369–385.

Sun, H., Lagarrigue, F., Wang, H., Fan, Z., Lopez-Ramirez, M.A., Chang, J.T., and Ginsberg, M.H. (2021). Distinct integrin activation pathways for effector and regulatory T cell trafficking and function. The Journal of experimental medicine 218.

Sun, H., Liu, J., Zheng, Y., Pan, Y., Zhang, K., and Chen, J. (2014). Distinct chemokine signaling regulates integrin ligand specificity to dictate tissue-specific lymphocyte homing. Developmental cell 30, 61–70.

Tadokoro, S., Shattil, S.J., Eto, K., Tai, V., Liddington, R.C., de Pereda, J.M., Ginsberg, M.H., and Calderwood, D.A. (2003). Talin binding to integrin beta tails: a final common step in integrin activation. Science 302, 103–106.

Takagi, J., Petre, B.M., Walz, T., and Springer, T.A. (2002). Global conformational rearrangements in integrin extracellular domains in outside-in and inside-out signaling. Cell 110, 599–511.

Takahashi, M., Dillon, T.J., Liu, C., Kariya, Y., Wang, Z., and Stork, P.J. (2013). Protein kinase A-dependent phosphorylation of Rap1 regulates its membrane localization and cell migration. J Biol Chem 288, 27712–27723.

Takahashi, M., Li, Y., Dillon, T.J., and Stork, P.J. (2017). Phosphorylation of Rap1 by cAMP-dependent Protein Kinase (PKA) Creates a Binding Site for KSR to Sustain ERK Activation by cAMP. The Journal of biological chemistry 292, 1449–1461.

Tkachenko, E., Sabouri-Ghomi, M., Pertz, O., Kim, C., Gutierrez, E., Machacek, M., Groisman, A., Danuser, G., and Ginsberg, M.H. (2011). Protein kinase A governs a RhoA-RhoGDI protrusion-retraction pacemaker in migrating cells. Nat Cell Biol 13, 660–667.

von Andrian, U.H., and Engelhardt, B. (2003). Alpha4 integrins as therapeutic targets in autoimmune disease. N Engl J Med 348, 68–72.

Wang, T.F., Lai, N.S., Huang, K.Y., Huang, H.L., Lu, M.C., Lin, Y.S., Chen, C.Y., Liu, S.Q., Lin, T.H., and Huang, H.B. (2012). Identification and characterization of the actin-binding motif of phostensin. International journal of molecular sciences 13, 15967–15982.

Wegener, K.L., Partridge, A., Han, J., Pickford, A.R., Liddington, R.C., Ginsberg, M.H., and Campbell, I.D. (2007). Structural basis of integrin activation by talin. Cell 128, 171.

Xiong, J.P., Stehle, T., Diefenbach, B., Zhang, R., Dunker, R., Scott, D.L., Joachimiak, A., Goodman, S.L., and Arnaout, M.A. (2001). Crystal structure of the extracellular segment of integrin alpha Vbeta3. Science 294, 339–345.

Ye, F., Hu, G., Taylor, D., Ratnikov, B., Bobkov, A.A., McLean, M.A., Sligar, S.G., Taylor, K.A., and Ginsberg, M.H. (2010). Recreation of the terminal events in physiological integrin activation. The Journal of cell biology 188, 157–173.

Zhang, Y., Kim, T.H., and Niswander, L. (2012). Phactr4 regulates directional migration of enteric neural crest through PP1, integrin signaling, and cofilin activity. Genes & development 26, 69–81.

